# Impaired context disambiguation as a key determinant of post-traumatic psychopathology: A neurocomputational account

**DOI:** 10.1101/2025.07.07.663473

**Authors:** Adam Linson, Heidi Anderson, Sarah Markham, Karl Friston, Michael Moutoussis

## Abstract

We combine theoretical biology and systems neuroscience to relate the mechanisms of post-traumatic stress disorder (PTSD) models to biopsychosocial complexity. Moving beyond fear conditioning, extinction and habits formed by associative learning, we consider conserved neuromodulatory mechanisms related to harm expectation, defensive mobilisation, and context (in)sensitivity. Our account of harm-mitigating, neurobehavioural responses furnishes a mechanistic understanding of maladaptive behaviour and impaired belief-updating, which are often underspecified in conceptual analyses. Grounded in neurobiological evidence, we specify PTSD-related impairment in terms of Bayesian belief-updating and establish an adaptive/maladaptive boundary using active inference simulation studies. The ensuing model plausibly illustrates PTSD pathogenesis, which remains understudied relative to aetiology. We locate pathogenesis in maladaptive precision following trauma, characterised by a distortion of ideal Bayesian updating via aberrant credit assignment. Future scientific and clinical studies can test the validity of these insights, which could underwrite our understanding of the phenomenology, diagnostics and ultimately treatment intervention in post-traumatic disorders.

## Introduction

During traumatic experiences, the imperative to mitigate against severe harm requires an intense mobilisation of biological subsystems, an ‘emergency response mode’ [1,2]. However, multiple factors involved in the traumatic experience, traumatic memory and the accompanying experience-dependent plasticity can lead this emergency response mode to become persistent, that is, maladaptive. Here, we focus on modelling post-traumatic pathogenesis as the process by which a trauma schema (or ‘mode’) becomes persistent and, in turn, causes a wide range of post-traumatic stress syndromes (PTSS), including those related to mental imagery (e.g. ‘flashbacks’). The modelling also illustrates the process of returning to a pre-traumatic response mode, to suggest how PTSS treatment interventions could target the root cause of symptoms. In contrast to reductionist accounts that can lead to over-medicalisation, this computational approach is consistent with a biopsychosocial and environmental understanding of post-traumatic disorders.

The present work models conserved neuromodulatory and sensorimotor mechanisms related to harm expectation, defensive mobilisation [3,4], and learning of causes and consequences (model-based learning) [5,6]. The model remains compatible with multiple accounts of fear and extinction [7–10], without relying on these constructs, by shifting the focus to the general construct of harm. Notionally harmful administrations of pain, electric shock, restraint, etc., are common to experimental paradigms of fear conditioning and extinction. Perception of noxious stimuli, expectation, and mitigation are necessarily evolutionarily conserved, as harm negatively impacts the survival of individuals and species.

The notion of expectation has been operationalised in research using a range of comparable theories including Hebbian learning, spike-timing dependent plasticity (STDP), predictive coding, and embodied cognition frameworks broadly known under ‘Bayesian brain’ or ‘predictive processing’ descriptions, sometimes specified as active inference (applied here). These theoretical approaches relate to methodologies for understanding and measuring correspondences between neural substrates and behaviour. Computational psychiatry commonly incorporates these methodologies [11], found in an emerging body of research on understanding traumatic experience and PTSS [1,12–14].

Neurobehavioural response patterns necessary for survival form the basis of our PTSD modelling. The sequence of agent-environment interactions engaging these neural mechanisms are: (1) learning about the sources of harm in multiple contexts; (2) learning when harm is probable; (3) making a (fallible) judgement about the likelihood of imminent harm; and (4) responding defensively only when harm is likely. Grounded in the anticipation of harm under varied conditions, the model remains consistent with experimental studies of fear conditioning and extinction in animal models [9], in addition to observational studies and clinical trials of human PTSD [15]. Nevertheless, we argue that probabilistic learning about contexts is central to understanding post-traumatic impairments [5,16], beyond associative (model-free) mechanisms often used to describe traumatic learning.

Beyond context learning, we draw on the neuroscience of visual attention, multimodal perception, memory, and sleep, and combine them with neuromodulatory systems neuroscience. On this basis, we articulate a general neuropsychological and neurobehavioral account of the putative pathogenesis of disorders related to traumatic stress, including PTSD and CPTSD. The account is critically evaluated by consulting Experts by Lived Experience (ELEs) of trauma. We conclude that the resulting neurocomputational model captures relevant biopsychosocial and environmental dynamics for understanding – and supporting future treatment of – these disorders.

A brief formulation of our analysis follows, linked further below to background literature and modelling. Throughout, we underscore contiguities between realistic experiential narratives and experimental paradigms (e.g. psychophysics). This approach extends our theoretical account beyond neurobiology and facilitates future empirical testing [17].

From the standpoint of an individual in a new situation, there may be ambiguity as to whether the surrounding context is one in which harm is likely (harmful context) or unlikely (harmless context). Ordinarily, determinations of context (harmful vs. harmless) are made by gathering sensory evidence to reach a decision. The belief that I am in one context and not the other should ideally persist to some extent as a (contextual) prior belief, but should also decay, in order to strike a balance between (a) maintaining the belief that I am still in the same context and (b) re-assessing whether the context has changed.

Our theory describes PTSD pathogenesis as a process by which the balance between harmless and harmful contextual priors is lost. Here, the prior belief in a harmful context grows (in precision) beyond a critical tipping point. Once reached, the aforementioned decay rate is insufficient for rebalancing with contextual priors for harmless context.

Following traumatic exposure to a harmful context, the imbalance occurs depending on resilience or vulnerability. For a resilient phenotype, the learning rate for harmful context is sufficient to preclude an imbalance between contextual priors (harmful vs. harmless). For a vulnerable phenotype, if prior beliefs about the presence of a harmful context become unduly strong, then opportunities for re-assessing context will be continually foregone. The foregoing of contextual re-assessment will extend to harmless contexts, due to the persistent (false) belief they are harmful. Moreover, without re-assessment, a (false) conclusion that the context is harmful will in turn be (mis)counted as new (false) evidence that the context is indeed harmful – thus continually strengthening the false contextual belief in a positive feedback loop, a ‘vicious circle’. This miscounting can be described as part of a confirmation bias in perceptual decision-making modelled by hierarchical approximate inference [18].

If PTSD pathogenesis occurs, treatment interventions provide and, crucially, remind one of evidence that one is in a harmless context, rather than a harmful one as believed *a priori*. For the treatment to succeed, one must accumulate enough evidence to update contextual priors. Only then can one seek novel situations and determine their true context, rather than relying on the harmful prior.

Next we review relevant literature to shape a common cognitive psychological framing of PTSD phenomenology. As a key example, we identify contextual interactions with the process widely described as Visual Working Memory (VWM). We then introduce a mechanistic account of neuromodulatory function in PTSD-related phenomena, and show how the neuroscience in question relates to our modelling approach. Finally, we illustrate PTSD pathogenesis and treatment in model-based simulation studies.

## Models and methods

### Post-traumatic cognitive psychology and predictive processing

#### The importance of context in remembering and acting

In everyday life, it is important to learn from a given encounter with a hostile person or adverse situation, to help mitigate potential harm in subsequent encounters. The canonical psychological account of this experiential process has proceeded in a bottom-up fashion, from perception to VWM to short-term and/or long-term memory (STM and LTM), and top-down from static memory to active recall and/or recognition. On the other hand, hierarchical predictive processing accounts cast this process as involving a multiple feedback and feedforward re-entrant loops, throughout the brain. Predictive processing speaks to a correspondence between the psychological constructs and interacting yet discrete neuronal populations implementing active sensing and learning [19–21]. Thus, our predictive processing modelling (specifically, active inference) is compatible with both psychological and neurophysiological explanatory frameworks [22,23]. We thus bring together wide-ranging evidence on PTSD, including cognitive psychology, biological psychiatry and related work on unimpaired mechanisms and processes, going beyond superficial connections to ground their fundamental integration.

When an individual has competing partially-activated event memories, as may be the case when happening upon a vaguely familiar scene, a sensory stimulus (e.g. a specific scent) associated with one memory but not the others can facilitate a (non-pathological) ‘flashback’ of the corresponding event [24,25]. This matches the famous Proustian phenomenology of having a madeleine with tea and being mentally transported to a vivid childhood scene as if it were present again, a re-experiencing. In such cases, a ‘winner takes all’ convergence on a decision amounts to an inference: inhibiting ruled out event memories (competing dynamic representations) and disinhibiting the selected one [26].

In cognitive psychological and neurobehavioural studies of how the past informs the present, experiments are often designed to mimic a naturalistic progression that involves event-based experience, memory, generalisation and recall. These can amount to learning that a policy (e.g. pulling a lever) has a desirable (familiar or preferred) result in one context and an undesirable (ambiguous or surprising) result in another. Learning about the policy in the context can allow a contextual cue to elicit the learned context-relevant propensities (e.g. approach vs. avoid). Human and nonhuman primate studies and computational simulations indicate that such task and context information is differentially encoded in memory [27–29].

Hippocampal dynamics are implicated in both generalised task guidance and episodic recall [19,25,30]. The potential for interplay between distinct generalised task-oriented behaviour and context-specific episodic memories offers an example of neural machinery impaired in PTSD. Normally, if you move heavy boxes, you may learn a generalisable task-relevant strategy for how to lift, hold and carry. Months later, when moving furniture, an association between the two contexts can help guide your behaviour. At the same time, the association may also lead to recalling specific episodic moments (‘flashes’) from moving boxes [31]. The contextual similarity between both events allows for generalising the task behaviour, while the contextual break between them facilitates their ability to co-exist in distinct event memories [30,32,33]. In the computational neuroanatomy of active inference modelling, context is encoded in a specified **D** vector, and associated with neuronal representations in the hippocampus [34,35]. With respect to hippocampal interplay between task-relevant and episodic context, modelling (below) shows how an inferred harmful context can elicit defensive behaviour and simultaneously promote introspective attention to event memory.

#### Cued attention and post-traumatic pathology

Non-pathological ‘flashes’ can have a similar genesis as clinical post-traumatic ‘flashbacks’. If a quotidian but non-ubiquitous object is part of a traumatic event, say, a readily available jacket worn by an aggressor in a hostile encounter, there is potential for a later encounter with someone else wearing the same type of jacket to elicit a traumatic memory.

Building on this example, a *resilient* individual who experienced the traumatic event may form an initial trauma association with the jacket, co-existing with other memories of seeing the same type of jacket on others. The association may be weakened over time, as the jacket cue is neuronally linked across distinct event boundaries. A repeat encounter with the cue will thus be insufficient to disambiguate co-activated memories, so that trauma memories do not become vivid or ‘capture’ attention. Attention will be easily deployed to the sensory periphery to accumulate evidence about present external surroundings. It is unlikely the present situation will be confused with the past event.

An illustration is given in Fig. 1, with a triangle standing in for the jacket. In Fig. 1 panel A, the triangle is in common to both circles, depicting event memories as guiding contexts. Seeing the triangle evokes both events, with a square or cross context, respectively. Deploying attention to the sensory periphery (Fig. 1, panel 4A) resolves the uncertainty about the present context.

**FIGURE 1.**
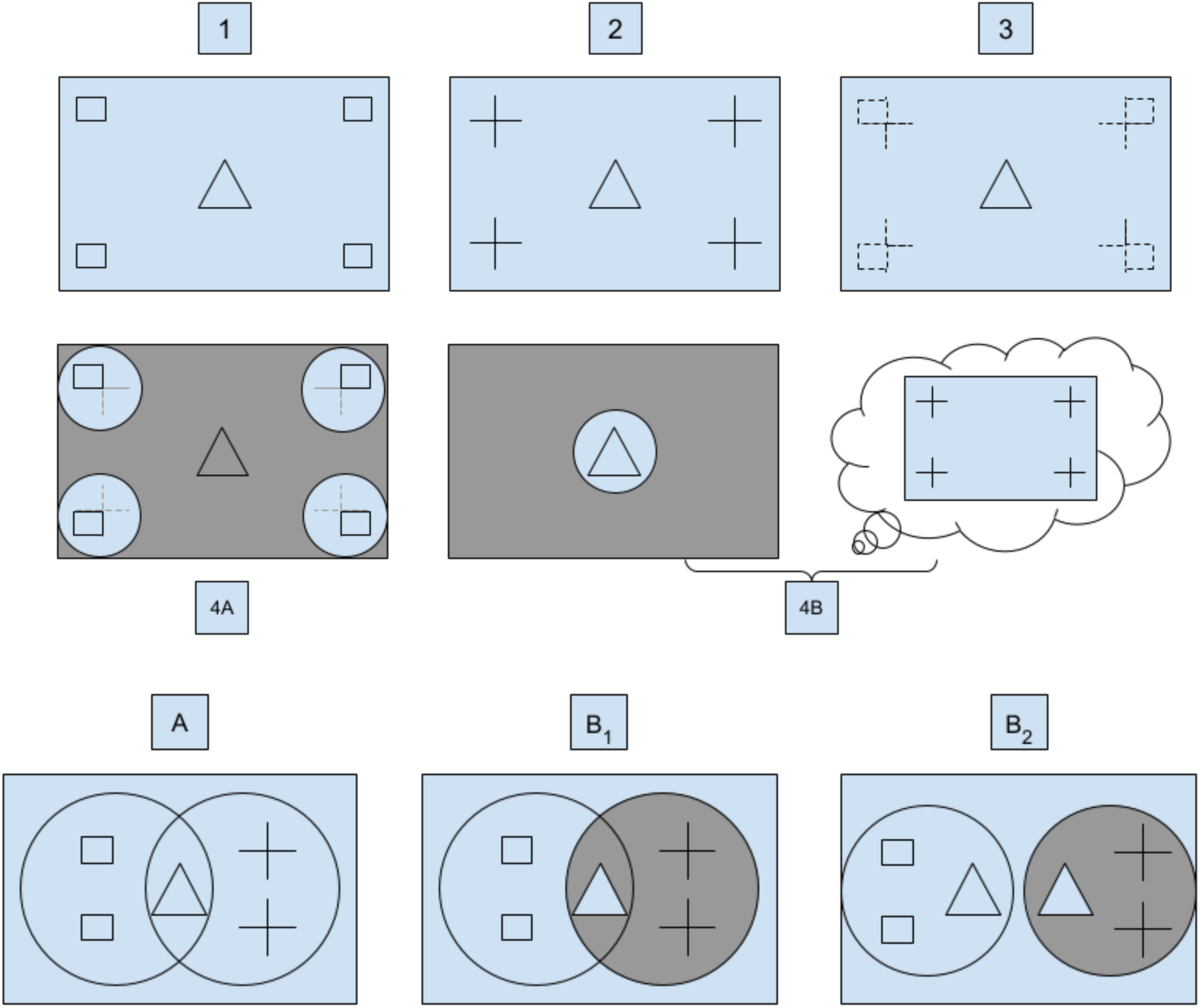
Two panels with a common element (△) and different backgrounds (□ vs. +) are labelled 1 and 2, respectively. After sufficient training to gain familiarity with panels 1 and 2, seeing the triangle in panel 3 suggests two alternative possibilities: it ambiguously matches panel 1 or 2. For this figure, it is stipulated that the veridical match is between panels 1 and 3. Even if panel 3 is predicted to match panel 2, attention can be directed to the periphery for sensory sampling (illustrated in panel 4A), to accumulate evidence about the veridical context. Once sensory evidence at the periphery is accumulated, it resolves the ambiguity of the background scene, leading to the inference that panel 3 context matches panel 1. A combination of sensory attenuation at the periphery with sensory focus on foreground, and high precision on recall of panel 2 – illustrated in panel 4B – can inform context disambiguation in favour of a false inference that panel 3 matches panel 2. See further below for a description of panels A and B_1-2_.

In a similar situation, a *vulnerable* individual may find the jacket cue capable of fully activating the trauma recollection, at the expense of any competing associations (Fig. 1, filled-in circle in panel B_1_; the contrast to B_2_ is explained below). This ‘winner takes all’ activation would predominate in VWM, increasing recall vividness and associated affective and motor responses, intermingled with present perception, causing a flashback (Fig. 1, panel 4B). The present salient object (triangle/jacket) interacts with the recalled traumatic context, mutually reinforcing the belief that the past event is present – in other words, a re-experiencing. On this understanding, an active inference account predicts increased activation of traumatic context representation. In contrast, dual representation theory (DRT) predicts increased weight on a bottom-up sensory pathway with an absence of top-down modulation by context [36]. It should be possible to distinguish experimentally between these predictions.

The attentional processes of the vulnerable person are likely to involve suppressed (attenuated) attention to peripheral sensory evidence that is potentially disambiguating. Due to this policy, one would suppress awareness and learning of the present, veridical context. This lack of learning reinforces a pathological persistent belief of being in the harmful context, given the cue (illustrated in our simulations below). Clinically, re-experiencing is self-categorized as injurious, and deliberate experiential avoidance further impairs learning. But even without deliberate experiential avoidance, the re-experiencing dynamics that limit peripheral awareness ultimately block inference and learning about benign causes of sensations (counterfactual blocking) [11]. In contrast, if counterfactual evaluation of context is not blocked, it becomes possible to recognise the veridical context. Even minor indications that I am not in a harmful context, if sustained (e.g. with adequate social support [37]), can facilitate a gradual updating and rebalancing of contextual priors that restores appropriate context-sensitivity. This rebalancing can be understood in terms of experience-dependent neuroplasticity. Restored context-sensitivity would be adaptive, as opposed to an ‘extinguished fear’, which could be maladaptive.

Recent findings show that stressful initial recall (as could be expected in a post-traumatic period) can prevent the formation of new engrams of the event memory [38]. Without typical post-consolidation processes, subsequent trauma recall can reactivate the original experience-encoded memory trace, consistent with traumatic re-experiencing, in contrast to non-traumatic unpleasant memory recall [39]. Fig. 1, panels B_1-2_, illustrate two ways in which the strength of an adverse event memory (dark grey circles) can furnish non-veridical context in a pathological flashback. (The two alternatives are not distinguished in our computational model, but future work will explore the relevant computational differences and their putative hippocampal basis.)

#### Traumatic memory and sleep

Our analysis and modelling provide fresh insight into the finding that dysregulated sleep following traumatic experience predicts the development of PTSD [40]. As described above, stressful recall results in the maintenance of a wakeful trauma experience memory trace that is reactivated on recall but not re-encoded [38]. There is another opportunity for re-encoding during sleep. Successful re-encoding facilitates a closed event boundary with selected associations neuronally linked across events (for instance, the jacket from the above example; see Fig. 1 panel A and B_1_) [33]. However, if following a traumatic experience, subsequent sleep is disrupted, then the original traumatic memory trace may continue to be maintained (Fig. 1 panel B_2_) [41].

In a healthy scenario, memories are repeatedly re-encoded in shifting sets of synaptic connections that are activated when recalling (’retrieving’) the original event [42]. This healthy progression depends upon retrieval while psychologically safe and during sufficient NREM sleep, both of which are challenged following acute or extended traumatic experience [38,40,43]. For example, childhood adverse experiences (CAE) can lead to neuronal ensembles encoding their memory under conditions that impede the healthy re-encoding process; this can in turn lead to the preferential re-activation of these ensembles during related adult stress [44], suggesting a contributing factor to CPTSD.

The neuronal ensemble initially encoded during an event overlaps with ensembles of re-encodings across repeated retrieval. In health, the re-encoded ensemble progressively ceases to overlap with the original one. This process can be understood formally as information compression or model reduction which minimises representational complexity (and maximises evidence for neuronal schemas in the absence of sensory input) [45], differentially driven by time and experiential learning factors [46]. These factors are captured indirectly (abstractly) by our models below: the time factor can be understood as a decay rate which is potentially too low in a person vulnerable to post-traumatic syndromes. If there is insufficient decay over time, alternative synaptic weights can be strengthened by learning (e.g. in treatment intervention). Such learning could reduce excessive re-activation of the original ensemble, which may subsequently support its healthy re-encoding.

A single nightmare or even a period of nightmares (e.g. following a bereavement) is not pathological, but may contribute to a period of intrusive cued re-activations in a manner suggested by our model. While our framing and modelling deliberately excludes fear and extinction, related work analyses nightmares in terms of fear extinction linked to ‘decontextualization’ and ‘recontextualization’ [47]. Fear extinction can be re-interpreted as follows. If there is a cue in common to multiple memories, an overly precise contextual prior for ‘harmful’ may preferentially activate a traumatic memory that gives rise to a nightmare. Weakening the prior (synaptic strength) may reduce cued re-activation of that memory, akin to decontextualisation of the cue. Strengthening a competing contextual prior for ‘harmless’ may then allow preferential activation of non-traumatic memories, akin to recontextualisation of the cue. We relate post-traumatic symptoms affecting sleep to neurochemistry in the next section.

### Neuromodulatory circuits and PTSD pathogenesis

Here we focus on functional neuroanatomy, to construct plausible hypotheses regarding the neural pathways that subserve the psychological, cognitive and physiological processes involved in our account. We initially draw connections between PTSS and a pontine neuromodulatory subsystem. We then provide a schema that brings this subsystem into correspondence with non-pathological and pathological neurobehavioral dynamics, and relate these to psychiatric diagnostic criteria. Finally, we elaborate a long-range circuit relevant to pathogenesis and treatment of PTSD.

The reticular activating system (RAS) in the brainstem, implicated in sleep-wake cycle regulation, is involved in a range of sleep disorders connected to psychiatric conditions [48–50]. Under typical functioning, excitatory cholinergic projections from the pedunculopontine tegmental nucleus (PPtg) to the locus coeruleus (LC) stimulate an inhibitory noradrenergic projection from the LC to the PPtg. In this PPtg-LC (excitation-inhibition) interaction, acetylcholine (ACh) and noradrenaline (NA) are mutually stabilising, resulting in a balance characteristic of attentive wakefulness.

At the initiation of sleep, the PPtg continues to emit ACh but temporarily ceases direct excitation of the LC via innervation pathways, resulting in a marked rise in ACh and diminishment of NA. If, however, this pontine ACh-NA subsystem is at an (aberrant) elevated tonic level, one would expect sleep to involve vivid trauma replay nightmares (elevated ACh) and gross body movements (elevated NA), both of which are characteristic of PTSD [41,50–53]. Known PTSS are depicted in Fig. 2, in relation to ACh and NA levels.

**FIGURE 2.**
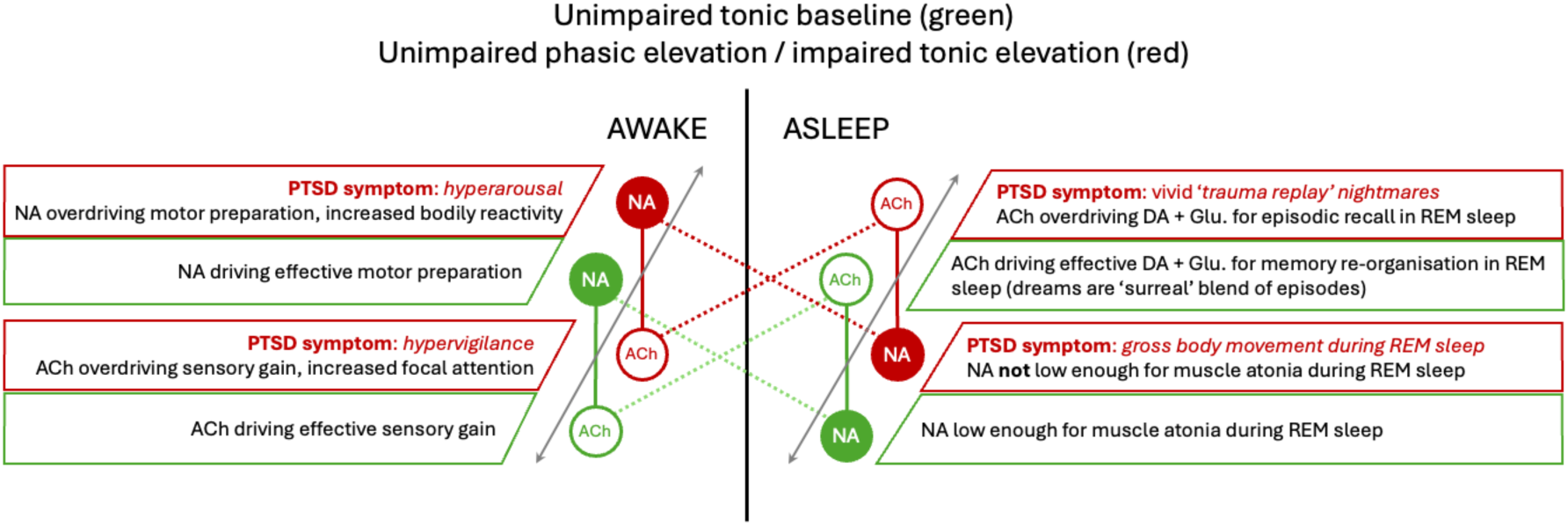
Across the central axis, the difference in relative ACh and NA levels in awake and asleep states is depicted (diagonal dotted lines). Green is used to indicate unimpaired tonic levels that raise to phasic levels (red) during a traumatic experience and return to unimpaired tonic levels (green) for a resilient individual (movement along the grey arrows). A vulnerable individual may remain at elevated phasic levels (red) such that these become tonic. The text boxes indicate corresponding consequences of ACh and NA when unelevated or elevated, including PTSD symptoms under maladaptive tonic elevation (awake and asleep).

The initial elevated tonic level (ahead of sleep) may be held over from an adaptive (i.e. situationally apt) phasic elevation during a traumatic event and its wakeful experience. During such an event, phasic elevation would aid harm mitigation efforts by increasing ACh for selective sensory gain and increasing NA for rapid behavioural response. Following such an event, phasic levels should fall ahead of healthy sleep, such that memory can be re-encoded and re-organised in relation to event boundaries.

Future reactions to trauma cues in new contexts could deploy attention to the sensory periphery to update beliefs about the veridical context (see Fig. 1, panels 4A and A). However, if the phasic increase has not subsided prior to sleep, then disrupted sleep can hinder memory re-encoding and re-organisation, resulting in stronger subsequent activation during traumatic recall. This overly strong reactivation can provide sufficient evidence for a false context, such that belief-updating about veridical conditions is hindered by peripheral sensory attenuation (see Fig. 1, panels 4B and B_1-2_). If the phasic elevation endures, it can effectively become a new elevated tonic baseline, with wakeful hypervigilance and hyperarousal. Fig. 3 illustrates how phasic elevation can become tonic elevation, and relates this to PTSD diagnostic criteria.

**FIGURE 3.**
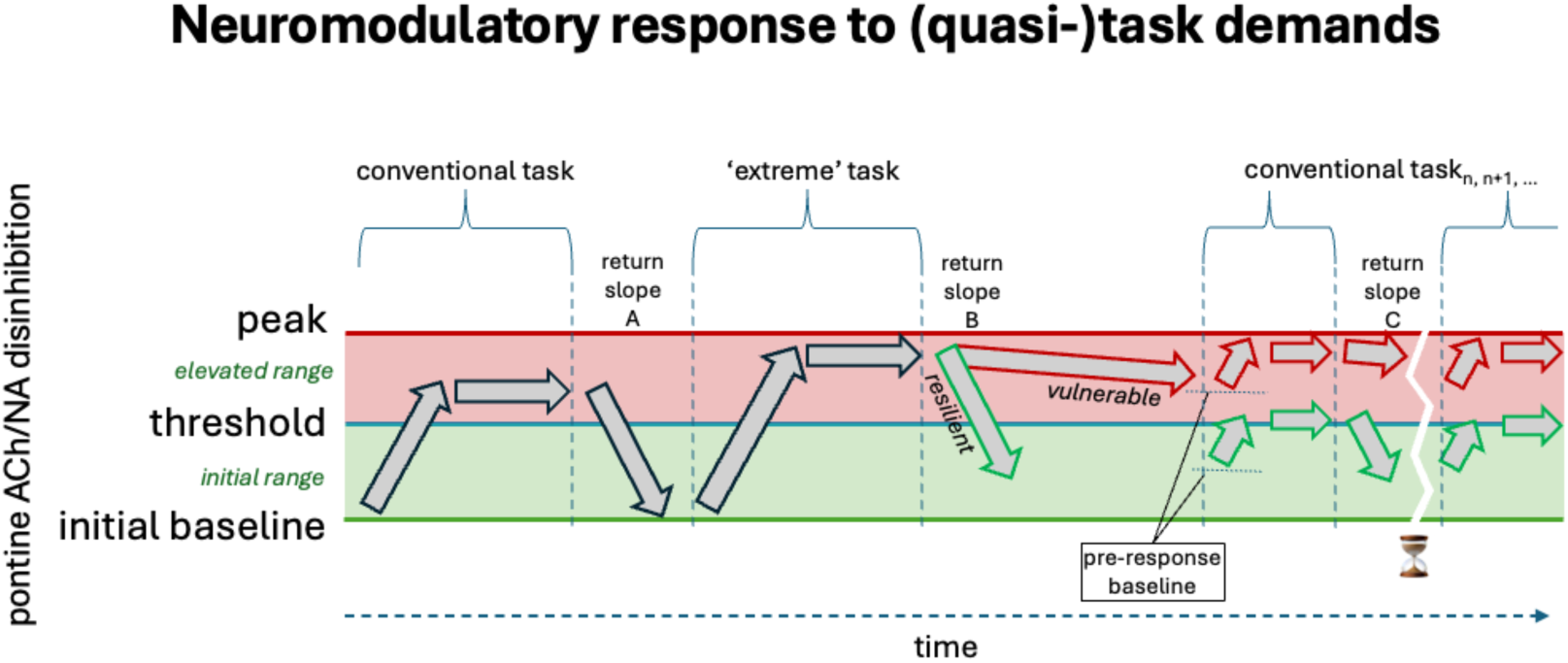
PTSS are natural, but their unwarranted persistence is pathological. Extreme task demands, as when life is at stake, elicit intense physiological responses, with physiological indicators peaking at high values (a time-course of sympathetic nervous system activation: increased heart rate (HR) and HR variability, blood pressure, breathing rate, adrenaline levels, vigorous motor preparation, etc.). How long it takes a defensive mobilisation to fully subside will depend on the elicited response (i) strength and (ii) the rate of response decay following a transition from a high to low probability of harm. An additional factor is (iii) the baseline immediately preceding the response, a local rather than global minimum (‘pre-response baseline’). If defences are already partly mobilised, as when on high alert, the distance between the pre-response baseline and peak mobilisation will be reduced (upper right arrows). For a diagnosis of post-traumatic stress disorder (PTSD), the prototype of its kind, symptoms must last “longer than a month” according to the DSM-5 but “at least several weeks” according to the ICD-11. From a computational point of view, the optimal decay rate should be determined by the persistence of high probability of harm. If, say, serious harm is likely to occur within three weeks, high alert should not stop within one week. Hence, we must not pathologise the initial occurrence of PTSD symptoms, but focus on how their nature increases the risk of unwarranted, maladaptive persistence. We postulate a critical relationship between a ‘return slope’ (B) and its intercepting a threshold separating extreme from ordinary arousal. This intercept characterises the transition out of PTSS – the time axis would be set differently in different diagnostic classifications and related to different neurobiological indicators (e.g. neurochemical and connectome properties [54]). Resilience can then be related to the ability to clearly infer that one no longer faces an increased likelihood of harm, and hence rapidly return to the psychophysiological baseline (i.e. ability to ‘bounce back’). An inappropriately sluggish decay of arousal would contribute to vulnerability to PTSS. Vulnerability factors may include genetic predispositions and lifespan events such as earlier psychological trauma, physical traumatic brain injury (TBI), ongoing adversity or comorbidity. The wide arrows indicate temporal increases and decreases in pontine acetylcholine (ACh) and noradrenaline (NA) levels (see Fig. 2 for corresponding symptoms and Fig. 4 for the circuit that controls these changes). Behaviourally, the responses relate to a naturalistic account of tasks (the term ‘task’ is used to link to experimental studies), divided into conventional tasks (e.g. securing food to satisfy hunger) and ‘extreme’ tasks (e.g. protecting oneself from serious harm). Following an extreme task (e.g. surviving a life-threatening encounter), defences may persist in a mobilised state (red, ‘elevated range’). Subsequent conventional tasks that would not have elicited a peak response when initiated at a low pre-response baseline may instead elicit a peak response, as they are continually initiated at a high pre-response baseline (upper right arrows). The high pre-response baseline will be maintained continuously given a slow decay rate (‘vulnerable’) and subsequent encounters with relevant sensory cues (described below). Thus, responses in conventional situations may frequently reach peak defensive mobilisation, constituting PTSS.

One ‘upstream’ reason the pontine ACh-NA subsystem may be disordered throughout sleep-wake cycles is reduced serotonergic (5HT) inhibition from the Dorsal Raphe Nucleus (DRN). The inhibition and disinhibition of this subsystem by the DRN emerges through a complex interplay of gamma-aminobutyric acid (GABA) and glutamate (Glu.) from the prefrontal cortex (PFC), limbic systems (LS), and basal ganglia (BG) [55]. Notably, altered GABA and Glu. levels and ratios have been found in PTSD, compared to unimpaired controls [56]. Taken together, these findings suggest a long-range circuit, by which forebrain interactions across the anterior cingulate cortex (ACC) can dampen DRN activity, thereby reducing the inhibitory influence of 5HT on pontine subsystems including the RAS, and especially the PPtg and LC. The PPtg and LC neuronal populations are the primary emitters of ACh and NA, respectively, with innervation throughout the brain in addition to reciprocal projections.

Excessive strength of the initial encoding of a traumatic experience is suggested by the increase in cholinergic stimulation from the PPtg to the Ventral Tegmental Area (VTA) and Substantia Nigra (SN). This ACh stimulation in turn stimulates a dopaminergic (DA) pathway to the forebrain (PFC, LS, and other parts of the BG). A DA surge is known to increase memory plasticity and encoding strength, especially during emotionally significant experience [19,57,58]. The full circuit is depicted in Fig. 4.

**FIGURE 4.**
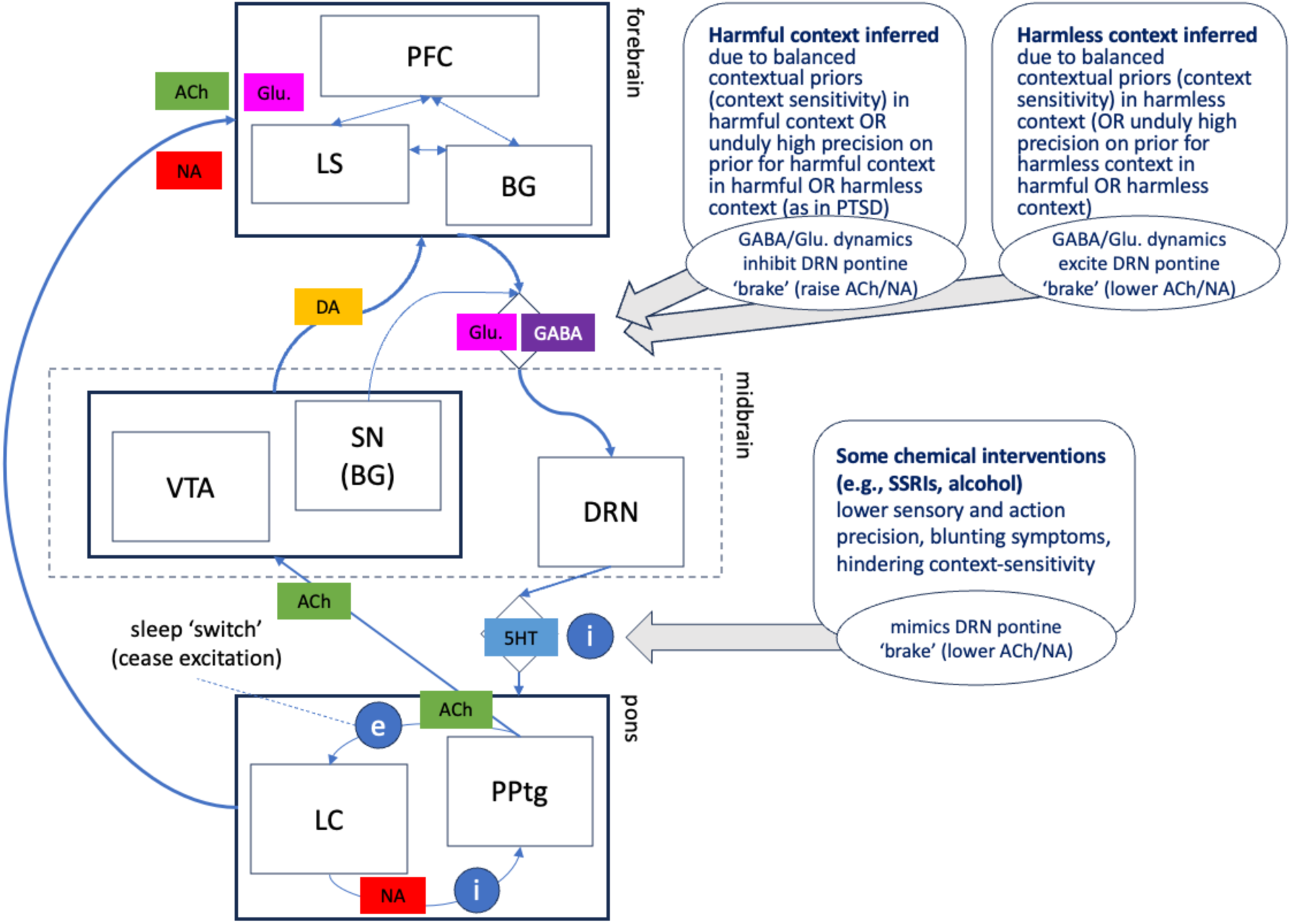
Neuromodulatory circuit. The neuromodulators noradrenaline (NA) and acetylcholine (ACh) are emitted in the pons. Cholinergic neuronal pathways from the pedunculopontine tegmental nucleus (PPtg) produce ACh and also have an excitatory connection to the locus coeruleus (LC). LC excitation produces NA that in turn has an inhibitory effect on the PPtg. This reciprocal relationship in the pons stabilises the ratio of NA and ACh perfusing the forebrain while awake. When asleep, the PPtg ceases excitation of the LC, allowing NA levels to drop and ACh to rise (‘sleep ‘switch’’).

With a stabilised ratio of NA to ACh, their baseline levels rise and fall jointly. The joint levels are modulated by the serotonergic projection from the midbrain dorsal raphe nucleus (DRN) to the pons. Increased serotonin (5HT) inhibits the pontine subsystem activity, thereby reducing its NA and ACh output. Conversely, decreased 5HT from the DRN disinhibits the pontine subsystem activity, allowing NA and ACh levels to rise.

The DRN is excited or inhibited by the interaction between glutamate (Glu.) and gammaaminobutyric acid (GABA), via glutamatergic and GABAergic projections from a richly interconnected network situated primarily in the forebrain. This forebrain network (with some outer nodes) is comprised of the prefrontal cortex (PFC), limbic system (LS), and the basal ganglia (BG), and also includes the anterior cingulate cortex (ACC) (unlabelled). When a harmless context is inferred via the forebrain network, the DRN ‘brake’ is applied; its serotonergic projections increase the 5HT input to the pons, inhibiting NA and ACh levels. When a harmful context is inferred, the DRN is inhibited (the ‘brake’ is released). In turn, the decreased 5HT input disinhibits the pontine subsystem, leading NA and ACh levels to rise.

PTSD pathogenesis can be understood in relation to the circuit diagram as follows. A current or recalled traumatic experience releases the DRN ‘brake’ to jointly raise ACh and NA for enhanced harm-mitigation (defensive response). A current harmful context will be strongly encoded with the potential to be easily cued. Later on, however, a resilient individual may experience cued traumatic recall, but with insufficient strength to resolve uncertainty about the present. This leads to peripheral sensory sampling, belief-updating about the veridical present, and subsequent applying of the DRN ‘brake’ to jointly lower ACh and NA (see Fig. 4).

In a vulnerable individual, the overly strong recall of a harmful context may cascade through jointly elevated ACh and NA, stimulated forebrain Glu. perfusion, and increased top-down excitement of VWM. Blending present visual sensation and recalled imagery in VWM (as described below) can lead to false contextual inference with rapidly evinced behaviours that the brain (falsely) infers as contextually apt. Thus, instead of an exploratory policy to resolve uncertainty about context, an exploitation policy is selected; namely, peripheral sensory attenuation and vigorous defence (fight / flight / freeze, etc.). This is consistent with a role for ACh enhancing attention to significant cues [59,60], amounting to hypervigilance (readily cuing harmful context), while NA simultaneously enhances motor responsivity [61], amounting to hyperarousal (see Fig. 2).

### Specifications for neurocomputational modelling

#### Context must mediate between perception and action

Stimulus-context associations extend to multimodal (polysensory) behavioural interaction with elements of the environment. Seeing an approaching object that looks as if it will soon hit me may coactivate the tactile expectation of being hit [3,62]. Visual recognition is not limited to sensory expectation, but also activates schemata that include overt behaviours embedded in the context evoked. For example, in a park, a defensive posture may be apt to protect oneself from being harmed by an approaching ball, initially seen at a distance [4,63]. Yet an equivalent visual sensation may cause an inapt defensive ‘flinch’ response during an immersive sports videogame when there will be no tactile impact. This scenario also describes the unimpaired ‘machinery’ that gives rise to PTSD when it is impaired.

A defensive reaction to current conditions can be understood as a consequence of prediction under a probabilistic (generative) model, as opposed to a stimulus-response account [22]. Under the active inference formulation, one is effectively working backwards from a future time point of avoiding (vs. experiencing) pain to the current time point of an approaching projectile. Inferences are made about the underlying (hidden) states that generate the sequence of impending sensations. On this basis, an action is selected that has the best chance of preventing a dispreferred outcome (being hit by a ball).

The context of the park vs. the videogame is relevant to the learned state transition sequence and resulting sensory outcome. When playing the videogame, what appears to be an incoming ball is a foreground cue that can lead to a false inference about the background context. If I (falsely) infer I am in the park, I expect the ball will hit me and cause a painful outcome. Thus, from a first-person perspective, a defensive flinch is apt (Bayes optimal), even though it is inapt to an observer who (correctly) infers I am playing a videogame. (Technically, this can be understood in terms of the complete class theorem [64,65]; namely, for every pair of behaviours and loss functions there are some prior beliefs that render my behaviour Bayes optimal. In the videogame example, these are empirical priors selected to rapidly infer the cause of sensations.)

Our focus in the present modelling is on prior beliefs about the context in which I am behaving. In this respect, we relate the above process of false inference about context to situationally inapt responses which, however, appear warranted in the presence of post-traumatic sensory attenuation. This is linked to our modelling through the notion of resource competition between traumatic memory and sensory present, where sensory sampling is attenuated due to high precision on a contextual prior.

#### Sensory attenuation is necessary to focus (cognitive) action

We argue that allocating attention is a decision central to both inferring the right context, and deciding upon the right actions. Below, we show *in silico* how inferring context and focusing attention feed off each other. Here, we introduce the principles of attentional control.

In a naturalistic, cluttered scene, individual object recognition may be diminished for most objects, especially when scanning a scene with unfocused attention. However, preferred (i.e. familiar or unsurprising) objects may remain undiminished despite clutter [66], which can explain a ‘pop-out’ effect when scanning a crowd and spotting a familiar face. This preferential recognition occurs in the ventral stream, which has been found to be impaired in PTSD, resulting in a form of sensory attenuation [67]. A puzzle about sensory attenuation in PTSD is how it might relate to hypervigilance, a form of sensory hyperactivity. This puzzle can be solved from the account we elucidate here.

Psychology and neuroimaging studies operationalise various load concepts (attentional, perceptual, cognitive, working memory) to focus on specific differences (e.g. visual pathways vs. attention networks). Yet the common ground among them is a resource model that can be mapped to neuronal populations with biophysical limits. For example, it may be that all neurons within a population are tuned to a single stimulus, or instead, a subset is tuned to (e.g.) another stimulus. From a technical perspective, this resource allocation is an emergent property of active inference in the following sense: the objective of active inference is to maximise the evidence for generative models of the lived world. Evidence (a.k.a. marginal likelihood) can be decomposed into accuracy and complexity. Therefore, model selection necessarily entails a minimisation of complexity, which can be read as minimising the degrees of freedom used to provide an accurate account of the sensorium [68–70]. Our current modelling does not aim to disentangle different aspects of resource allocation within active inference. For our purposes, it suffices to say that mitigating harm in a traumatic encounter commonly requires visuospatial attention and sensorimotor coordination; namely, efficient active vision [71–74].

Sensory working memory such as in the visual modality (VWM) is a useful construct exemplifying load-limited interactions related to visuospatial sensorimotor task demands. We treat VWM as a limited neuronal resource susceptible to ‘bottom-up’ (sensory driven) and ‘top-down’ (recalled imagery driven) excitation [20,75,76]. Other findings on visual pathways also bear, albeit indirectly, on our computational account. For instance, there is empirical support for reduced processing of peripheral visual stimuli under higher task demands, i.e. higher cognitive load; this includes the direct interaction between optical information and visual cortices [77–79]. In our account, the reduced influence of sensory evidence on VWM in some PTSS phenomena corresponds to a competing contribution of recalled mental imagery. We model this as an explore/exploit trade-off, mediated by the precision afforded to new observations vs. prior expectations.

#### From mental imagery to traumatic flashbacks

In PTSD, increased bottom-up gain has been found to co-exist with increased top-down gain [80]. Two connectivity pathways, from vision (bottom-up) and mental imagery (top-down) [75,76], can be understood as converging in VWM [20]. Even if a visual element in VWM is absent from a ‘bottom-up’ sampling of the scene, VWM can robustly maintain visual information ‘top-down’, as if the item were present [81]. Moreover, the top-down influence of mental imagery on VWM can affect and distort concurrent bottom-up visual perception [82].

In line with the scenario of a looming projectile above, VWM is not necessarily detached from motor behaviour. Mental imagery of a scene can even influence oculomotor movement, inducing saccades to fixation points that match saccades in perception of the same scene [83,84]. One can also pursue similar arguments to explain rapid eye movements during dreaming in REM sleep [45]. This coupling could explain why altered eye movements (as in EMDR therapy) would disrupt the vividness of mental imagery, and in turn reduce the magnitude of affective response to mental imagery of a traumatic experience [85].

A synthesis of the above evidence points to a possible explanation for ‘flashback’ phenomenology: high sensory precision or gain (modulated by ACh [51]) can facilitate visual attention to scene elements with enhanced salience (trauma reminders or ‘cues’) that are rapidly identified and rapidly elicit a trauma-context association. The traumatic context is characterised as one with a high probability of causes of harm (‘harmful context’) [16]. The on-going imperatives of harm-mitigation requires flexibly attending to ‘task-relevant’ stimuli; i.e. sampling the visual scene to maximise information gain (a.k.a. Bayesian surprise) [86–89]. This aspect of active vision follows the metaphor of an attentional ‘spotlight’ that must rapidly shift among momentary foreground elements. With each shift, the remaining background scene is relegated to a diminished ‘task-irrelevant’ periphery [79]. The restlessly roving spotlight would be indicative of hypervigilance, with increased sensory gain on a focal element and corresponding attenuation of the sensory periphery.

In terms of belief-updating, given the (false) belief one is in a harmful context that matches traumatic recall, a strong top-down influence of trauma scene imagery on VWM that is concurrent with bottom-up foreground perception can cause a distortive blend of the present scene and a past traumatic event. These imagery-based elements would continuously support misperception and misrecognition of the current scene, allowing attention to be maintained on a significant foreground element while hindering attention to – and subsequent belief updating about – the actual external context (Fig. 1, panel 4B). In turn, a behavioural response is elicited that would be appropriate to a repeat traumatic encounter, but is inappropriate to the present veridical conditions. This formulation of sensory attenuation theory thus provides a testable hypothesis for the neurocognitive machinery of ‘flashback’ phenomenology, in that cued traumatic recall is predicted to attenuate peripheral sensing more than focal sensing (e.g. in a target/distractor paradigm). The theory also provides a testable hypothesis related to another common PTSD symptom, the exaggerated startle response (ESR); namely, that alternatively valenced contexts for the same probe stimulus should differentially affect startle parameters for PTSD-impaired individuals to a lesser degree than those without [90].

### Model definition for simulation studies

We developed a neurocomputational model of interacting systems perspectives schematised above: the *neurocognition of trauma associations* including learning, memory, and context; the *phenomenology and symptoms of PTSD* and their relation to treatment intervention; and the *neurobehavioural dynamics of defensive mobilisation*, underpinned by a *neuromodulatory circuit*. We then performed model-based simulation studies that explain (i) natural waning of PTSS in resilient individuals under conditions of ordinary stress, (ii) persistence of such symptoms in predisposed individuals in similar conditions, (iii) recovery of context disambiguation through a learning-based intervention, and (iv) the limited role of symptom reduction through monoaminergic therapy on its own.

### Recurrent Attention-modulated Belief-updating of Context (ABC) model

The simulations of PTSD pathogenesis and treatment intervention below are based on a generative model (Fig. 5) that refines, extends and operationalises previous work [1,11,12]. They reproduce the pathogenesis of PTSD following a traumatic encounter, and its trajectory following a successful or unsuccessful intervention. The learning and behavioural dynamics of the system are driven by uncertainty resolution, implemented as free energy minimisation.

**FIGURE 5.**
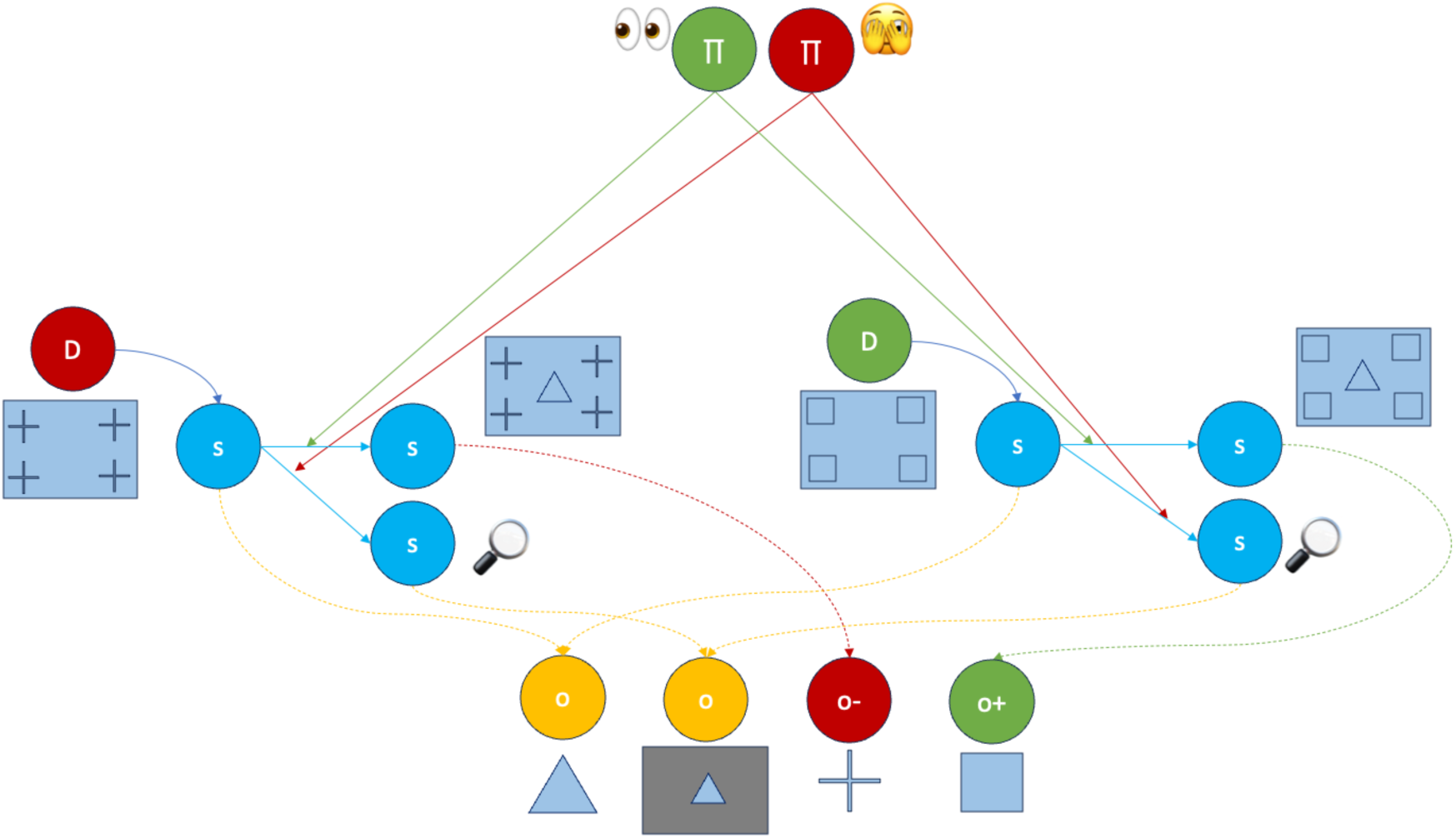
See main text for description.

In Fig. 5, the contextual priors (circles labelled **D**) condition the initial state (along arrow to blue circle), which generates a contextually ambiguous observation (yellow circle with free-standing triangle). During the initial traumatic experience, the system advances along the horizontal arrow from the initial state to the second state (blue circle with adjoining rectangle). The state sequence is controlled by an exploratory scene sampling policy (green circle with ‘two eyes’ pictogram) to resolve uncertainty about the context. This second state generates adverse (surprising) observations that can include a range of ‘harmful’ sensations (free-standing ‘+’), among them pain (e.g. being struck by shrapnel) and visual scenes that inflict moral injury (e.g. bearing witness to extreme suffering). The observation allows for a veridical contextual inference, updating the prior probability (red **D**) that one is in the harmful context.

Following the harmful traumatic experience, in a new harmless context, one begins in the initial state conditioned by the green **D**. This start state also generates a contextually ambiguous observation (free-standing triangle). In other words, this initial observation is insufficient to disambiguate between the two alternative start states (tracing back yellow arrows to the left or right). The exploratory policy progresses the state sequence along the horizontal line to generate context-specific observations (free-standing square). This disambiguates the two contexts, leading to a veridical inference of the harmless context.

However, if one begins in the ‘harmless’ initial state (conditioned by the green **D**), and the contextual prior of the past (traumatic) context is unduly precise, it is sufficient to resolve the uncertainty about the context, such that the exploratory policy is not selected (it is falsely believed to be uninformative due to overly informative prior beliefs). This results in false contextual inference and selection of the (peripheral) sensory attenuation policy (red circle with ‘partly covered eyes’ pictogram). We emphasise that this form of sensory attenuation is consistent with hypervigilance, amounting to rapid shifting of foreground focus without resolving contextual uncertainty.

From a first-person perspective, there is enough (misleading) information that one is in the harmful context, even though from a third-person perspective, one is (in fact) in the harmless context. Moreover, under the sensory attenuation policy, the foreground focus state (‘magnifying glass’ pictogram) generates the same observation irrespective of context (‘focal’ triangle in grey box). Like the initial observation, the focal observation is insufficient to disambiguate context, so the agent never learns they are in the harmless context.

As long as the sensory attenuation policy is selected, observations more consistent with harmless contexts will be too slow to overcome the bias towards crediting the harmful context with (unwarranted) evidence. In past work, we proposed that in vulnerable brains, Bayesian learning ‘breaks’ because of an impairment in the ability to accumulate evidence that a context is benign (counterfactual blocking). Our simulation studies suggest that the specific weights of learning and forgetting, through which the memories of encounters with harmful and harmless contexts accumulate, interact with learning history to stabilise the inappropriate beliefs (see below and Fig. 7).

#### Hierarchical prior dependent trauma re-experiencing

A second aspect of the generative model (Fig. 6) illustrates a secondary process that underpins ‘re-experiencing’ symptoms. It does not drive the simulation dynamics but rather emerges from policy selection under the core recurrent ABC model (Fig. 5). In this secondary process, high precision on a contextual prior can lead to a blend of sensation and imagery in VWM indicative of flashback phenomenology. A trauma-associated cue (triangle) can elicit multiple contexts (harmful vs. harmless) without sufficient information to disambiguate them. Under relatively balanced priors, the exploratory policy is selected (see Fig. 1 panels 4A and A). High sensory precision can be allocated to the periphery, even when there is sparse contextual information (see Fig. 1 panel 3). This allows for (bottom-up) sensory evidence accumulation about the present harmless context. Under highly imbalanced priors, with undue precision on the harmful context, the sensory attenuation policy is selected. This places low sensory precision on the periphery, allowing for VWM to be partly excited by recalled imagery (top-down), i.e. from the past harmful context (see Fig. 1 panels 4B, B_1-2_).

**FIGURE 6.**
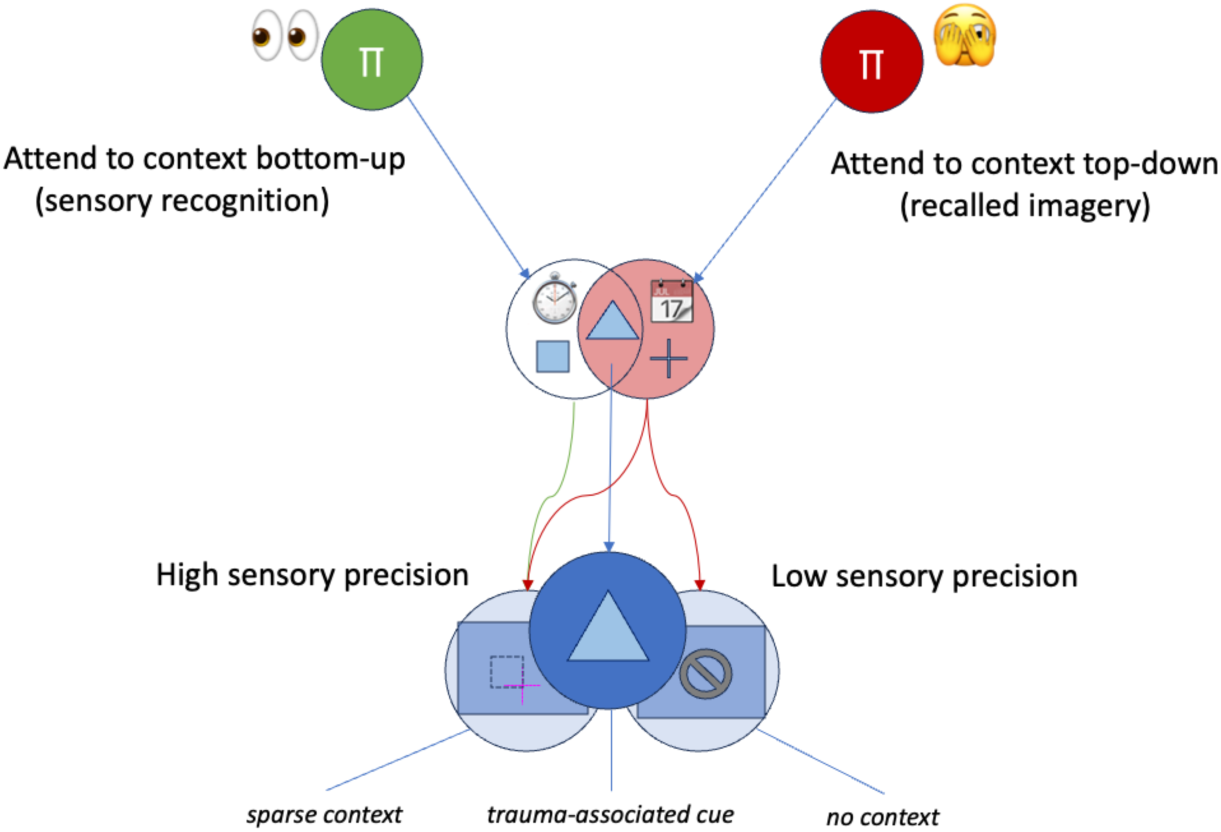
See main text for detailed explanation and correspondence to Fig. 1. Here, the stopwatch pictogram indicates the present (veridical) ‘harmless’ state and the calendar pictogram indicates a past ‘harmful’ state.

In other words, in the above scenario, a focal visual sensation would be congruent with focal mental imagery. This congruency, combined with peripheral sensory attenuation and strong contextual imagery, could meet a subjective ‘reality threshold’, amenable to experimental study in a reality testing paradigm [91]. To elaborate, a possible mechanism by which a percept meets the threshold for a ‘reality’ judgment involves the congruency of bottom-up sensory and top-down mental information. However, it may be possible for this mechanism to be undermined, given post-traumatic impairment and a trauma-associated focal sensory cue (Fig. 6). The focal element would be simultaneously present to the senses and present in recall, such that their congruency would strengthen a ‘reality’ judgment (i.e. updating the belief that present experience reflects external reality). At the same time, if peripheral sensory information is attenuated, the (potential) incongruency of available sensory information and traumatic recall of the peripheral scene would evade the reality testing mechanism altogether, as no incongruency would be registered. Without the incongruency that would ordinarily weaken a ‘reality’ judgment, traumatic recall blended with sensory perception of a harmless situation could be experienced with a false sense that a harmful situation is presently occurring in reality.

## Results

In Fig. 7, we see the evolution of contextual priors in a resilient (left) and vulnerable (right) system. Each system starts with two attractor basins that represent the initial priors **D**, with harmless (left) and harmful (right). Following a short sequence of harmful trials (that represent a traumatic experience) – followed by an extended sequence of randomly allocated harmful and harmless trials (50/50) – we see how the resilient system is able to garner veridical contextual information and update its priors. This leads to balanced priors that bias the system towards the exploratory policy for learning the true state of the world. In the vulnerable system, we see what amounts to a ‘vicious circle’ that characterises PTSD and related disorders: the sensory attenuation policy prevents learning about the harmless context, which biases the system towards the sensory attenuation policy, and so on, in a self-reinforcing loop.

**FIGURE 7.**
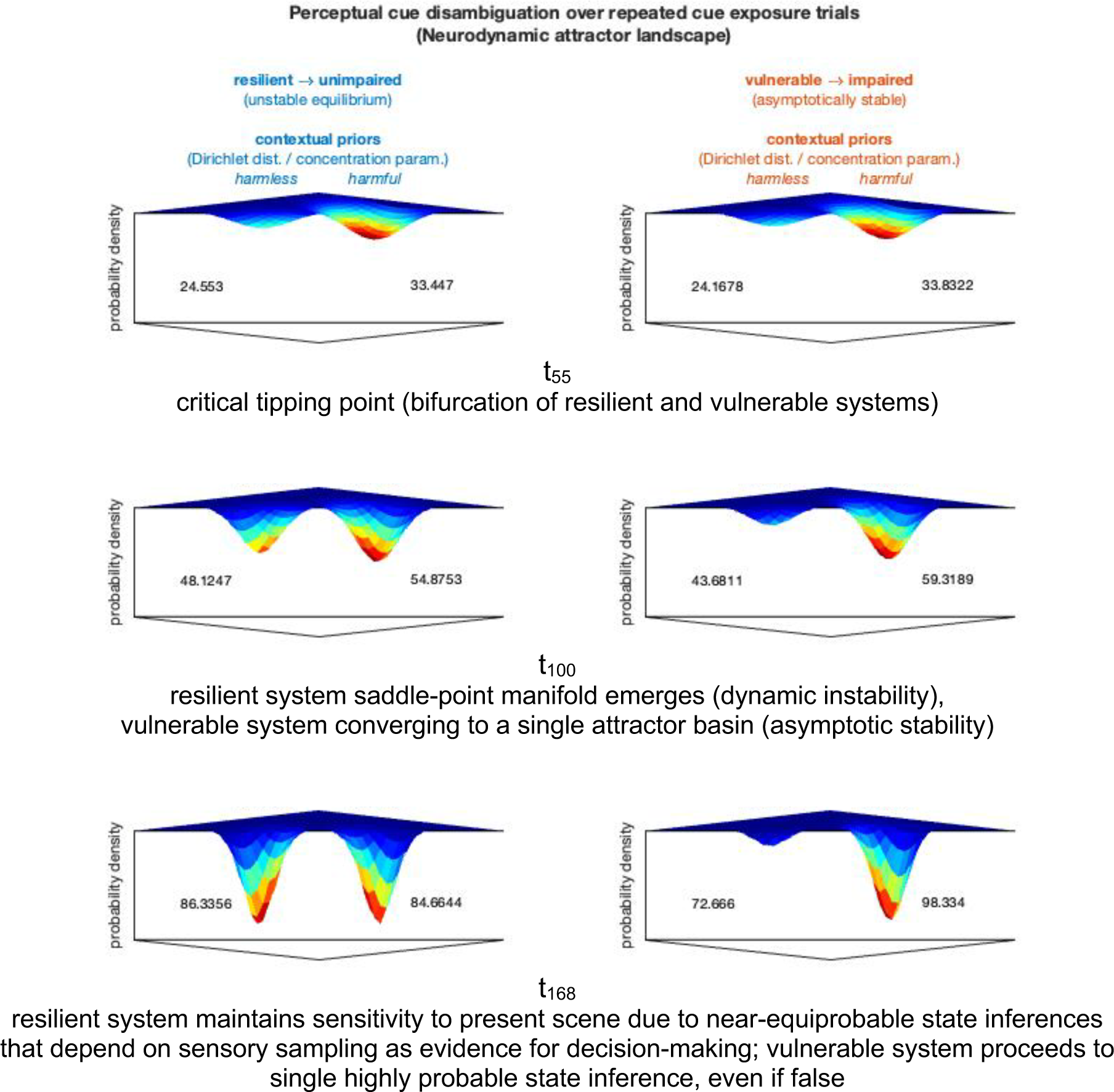
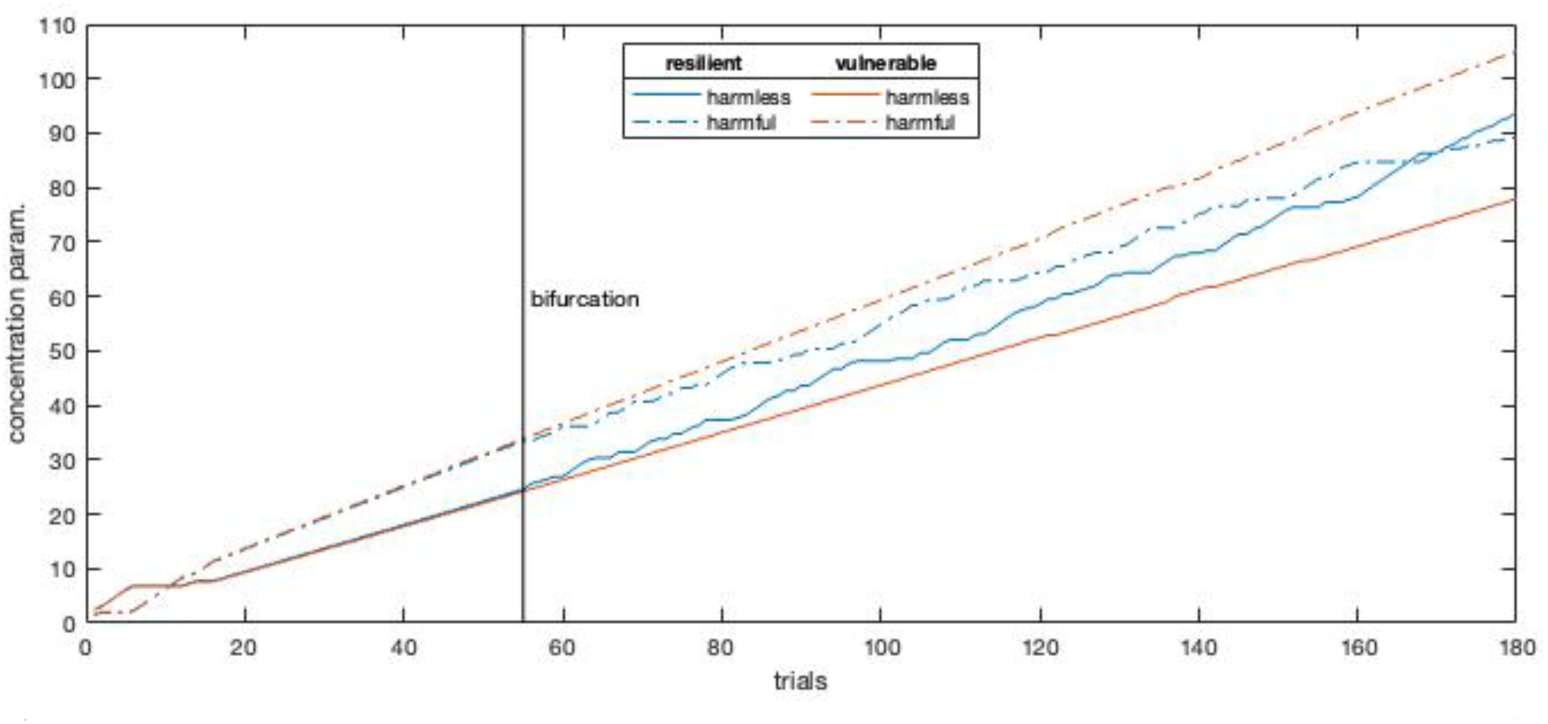
See main text for description of the attractor basins. Lower panel is an additional visualisation of the basins, where t55 is the first panel row above, labelled here as the bifurcation. The bifurcation arises from two different rates of harmful evidence accumulation.

The resulting landscapes at the conclusion of both the resilient and vulnerable processes are consistent with recent neural recordings of macaque perceptual decision-making [92]. In the macaque study, neurons accumulate evidence for two alternative visual fixation target options preceding a saccade-to-target action. When there is ambiguity between the targets, decision time is longer, as sensory evidence accumulates to resolve the ambiguity. When there is bias towards one option, the decision time to reach a commitment (action) is faster, and the bias functions as greater ‘pre-accumulated’, or notional, evidence.

Our case is not about alternative perceptual targets, but the interpretation is similar [93]. Following a cue (triangle) common to two contexts (squares and crosses), if there is insufficient evidence to resolve which context is present, then more evidence accumulation is required to resolve the ambiguity (higher decision time to commit). In our model, this is achieved by an additional timestep of peripheral sensory sampling, to accumulate contextual evidence for either squares or crosses (see Fig. 1, panels 4A and A). However, if there is a strong bias for one context (crosses), the contextual inference for crosses is made immediately, even if this amounts to committing to the incorrect decision when the true context is otherwise, i.e. false inference (see Fig. 1, panels 4B and B_1-2_).

In Fig. 7, the basin shape represents the contextual priors. The decision trajectory for resolving ambiguity must either continue to sensory sampling given the decision manifold under insufficient prior evidence (unstable equilibrium) or can forgo further sensory evidence due to a collapse into a single stable basin. The basins are illustrative data visualisations based on plots of the Dirichlet distribution **D**; they can be interpreted formally as the (free) energy landscape of a combined integration and decision-making network [94].

In Fig. 8, we show that a successful intervention is one that restores context-sensitivity to behaviour, linked to the rebalancing of prior probabilities. This success may be facilitated with or without psychopharmacological support. For example, SSRIs may be used therapeutically to improve the potential for new learning.

However, psychotropic drugs (including SSRIs, benzodiazepines, or excessive alcohol consumption) can also show illusory success when they blunt context-sensitivity in ways that reduce PTSS, without new learning. Interventions of this sort are modelled via changing the precision of the likelihood mappings (representing sensory precision) and transition priors (representing prior precision). Numerically, these interventions were simulated by lowering precision in the corresponding **A** and **B** tensors (under a generative model based upon a partially observed Markov decision process).

The disappearance of symptoms in the absence of context-sensitivity is insufficient to rebalance priors, leading symptomatic behaviour to return at equal or greater strength upon cessation of chemical intake. Conversely, cessation of (e.g.) a psychotherapeutic treatment that has facilitated an increase in context-sensitivity retains the potential to lead eventually to rebalanced priors. This contrast between successful and unsuccessful PTSD treatment closely matches randomised controlled trial (RCT) results [15].

Each diagram in **Figure 8a-e** can be interpreted as follows:

- The numbers along the bottom axis indicate the trial number in blocks of 60, comprising two alternative sequences of 180 trials (3 blocks of 60 each). The sequences are clarified in the captions beneath each block. (Summary: 7a depicts the initial 60 trials, with two alternative continuations of 2x 60-trial blocks, either to 7b-7c or 7d-7e. In other words, the 60 trials of 7a are in common to both sequences, and 7b and 7d begin with an alternative trial 61 of 180; 7c and 7e begin with an alternative trial 121 of 180.)
- The top row indicates whether it is a harmless (□) or harmful (+) trial.
- The four ‘internal’ rows (with greyscale boxes) indicate the policy selection for each trial. A trial is comprised of three time-steps: a start state, and two policy selections that advance to a middle and final state, respectively. (Fig. 5 depicts the pathways from the start state to the middle state, and the final time step allows the agent to remain in the middle state for the final state, or move to the alternative non-start state.) The four available policies are all the permutations and combinations of the two actions: 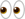=exploratory sensing (of the periphery) and 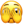=sensory attenuation (of the periphery). In the rows below, from top-to-bottom, the policy for two timesteps is: 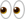-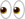(row 1), 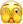-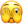(row 2), 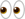-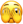(row 3), 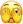-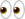(row 4). The grayscale within each trial column indicates the progressive resolution of uncertainty by belief-updating (within the policy space) until policy selection.

**Figure 8a.**
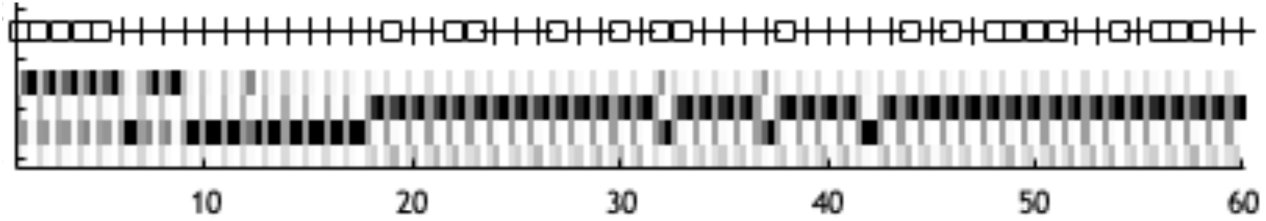
Initial sequence of trials. Trial 1-5: harmless condition (□), 6-16: harmful condition (+). 17-60: random. Without intervention, the sequence continues in the manner of trials 50-60 (i.e., repeated selection of sensory attenuation policy under any condition). The evolution of contextual priors for this continuation without intervention (from trials 1-60 above and 61-180) is shown in the vulnerable system in Fig. 7.

**Figure 8b.**
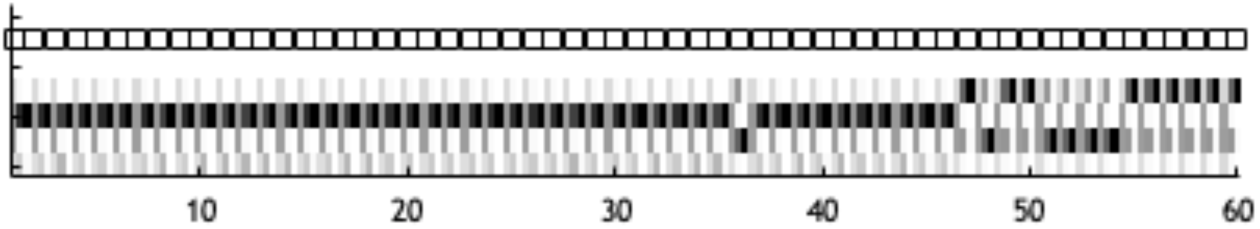
Simulated treatment option to restore context-sensitivity. Controlled exposure to harmless condition for 60 trials. Towards the end of the sequence, the exploratory policy begins to emerge and the contextual priors begin rebalancing. This can be understood in terms of Fig. 7, as a shift from a vulnerable individual (that continues along the vulnerable trajectory in the simulation with no intervention) to what resembles the resilient individual (who never develops the imbalanced priors in the simulation).

**Figure 8c.**
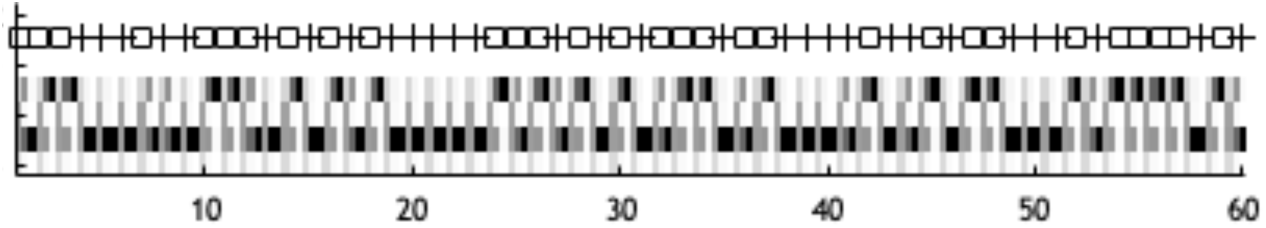
Following the treatment intervention simulated in Fig. 8b, the remaining trials continue at random (harmless or harmful), maintaining balanced priors and appropriate context-sensitive policy selection.

**Figure 8d.**
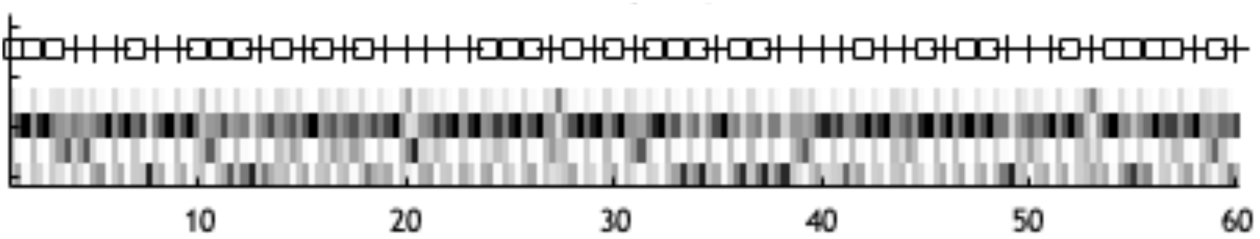
This sequence is an alternative progression following the 60 trials of Fig. 8a. It approximates the effects of (e.g.) SSRIs (selective serotonin reuptake inhibitors) or alcohol in lowering the precision of sensory information or action. At the same time, the trials are random to illustrate the absence of a controlled environment [95]. The kind of response vigour seen in Fig. 8a trials 50-60 is diminished here, indicated by more grayscale (low precision policy selection) and a wider range of selected policies, which can be interpreted as a reduction in PTSS.

**Figure 8e.**
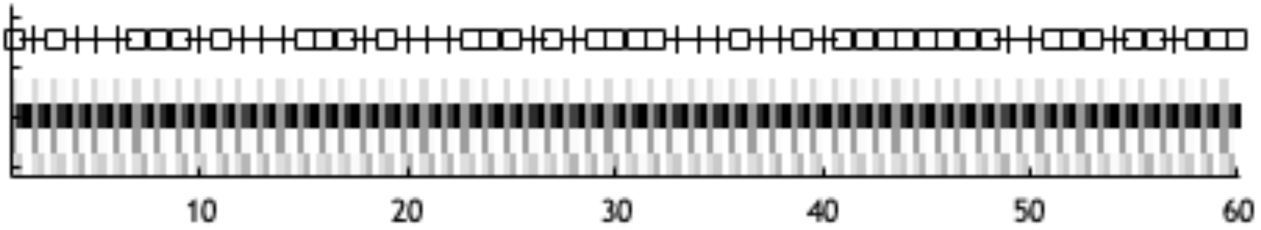
This sequence continues from Fig. 8d with the same random alternations of conditions, but with a restoration of the initial sensory and action precision from Fig. 8a, representing the cessation of chemical intake (e.g. SSRIs or alcohol). In the Fig. 8d trials, the contextual priors were not rebalanced, so here, the context-insensitivity governs the policy selection. As described in the main paper, an exploratory policy is falsely ruled out as unnecessary, given what is falsely taken as sufficient information about context from the prior. This means that even in the harmless trials (□), the sensory attenuation policy is selected. In line with the full analysis in the main paper, this amounts to a self-reinforcing neuromodulatory ‘vicious circle’ of defensive policy selection and counterfactual blocking.

**Figure 8f.**
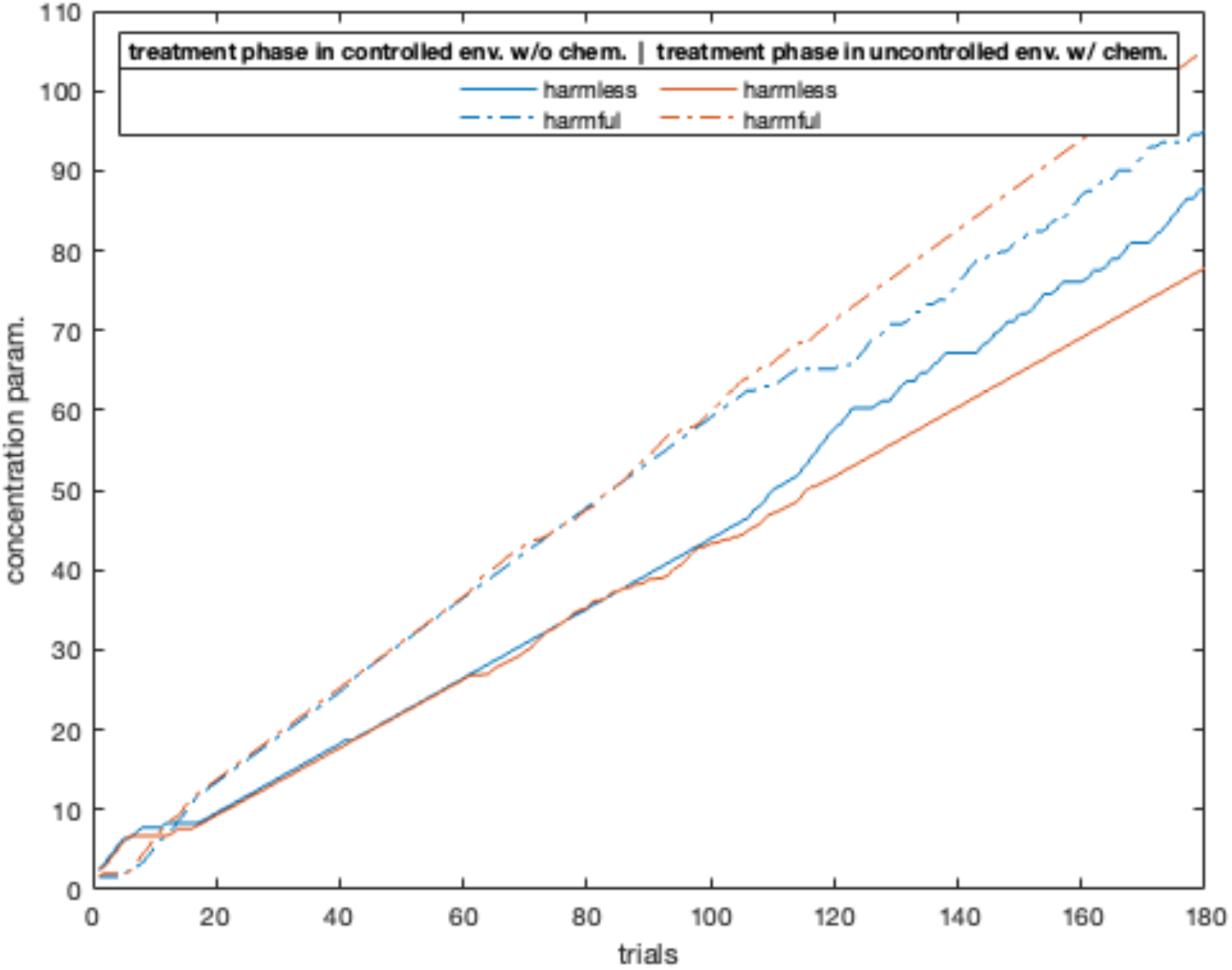
Evolution of contextual priors (vulnerable), with alternative interventions in trials 61-120. Gradually converging plot lines (blue) depict the trial sequence of Fig. 8a-**b-c**, with a simulated inter-mediary phase of no chemical intervention under a controlled environment (e.g., psychotherapeutic treatment). Diverging plot lines (orange) depict the trial sequence of Fig. 8a-**d-e**, with a simulated intermediary phase of chemical intervention (e.g. SSRIs or alcohol) under an uncontrolled environment (simulated by lowering precision in the **A** and **B** tensors).

### Discussion

The analysis and modelling presented here serves to formalise concepts that often remain underspecified in the narrative literature on PTSD and related psychopathology. In this work, context refers to elements of the sensory or cognitive periphery that provide evidence for the states of affairs prior to belief-updating. Belief-updating here refers to changing the neuronal representations of the likely properties of the world. Our simulations using the recurrent *Attention-modulated Belief-updating of Context* (ABC) model demonstrate the interdependence between inference and learning, and how biased inference can lead to aberrant learning and a failure to recognise changes in context. It is this failure we associate with the core psychopathology of PTSD.

Biased inference in this instance manifests as a failure of responding to epistemic affordances that leads to a failure to recognise the context has changed. This kind of failure sits on a critical tipping point, leading to a phenomenal description of the ensuing inferences in terms of bistability, that can be related to the notion of canalisation in psychopathology [96]. While attractor basins are used in different ways to indicate neuronal dynamics, here, we have shown that plasticity can create unstable basins that bias the system toward sensory evidence accumulation to resolve ambiguity about the causes of sensations. In contrast, a collapse or canalisation into a single basin can cause a trajectory initiated by a sensory cue to become inescapably bound by the basin, resolving uncertainty through precise – but false – beliefs about the causes of sensations. Crucially, this resolution is self-maintaining because it precludes epistemic foraging or information seeking, thereby instantiating a vicious circle of confirmation bias. Restoring plasticity can make the system amenable to new learning, thus rebalancing priors and restoring the collapsed basin. The saddle point between basins corresponds to a reinstatement of context-sensitivity.

In a context where harm is highly likely, such as a combat zone, it is adaptive to have elevated ACh and NA, to remain vigilant against possible harm, and rapidly mobilise a defensive response for mitigating harm. Essentially, it is context-sensitive to be in a heightened alert state, ‘better safe than sorry’. After a transition into a context where harm is unlikely, it does not mean harm is impossible. A home neighbourhood may be largely harmless, but occasionally harms can arise. Thus, context-sensitivity requires sensory disambiguation on a case-by-case basis, as a tap on the shoulder could be from a friend or foe. It is maladaptive to immediately respond to such a shoulder tap with a vigorous fight-or-flight response, as this would be unwarranted if it were a friend. Yet, it is also maladaptive to be ‘fearless’ and disregard learning whether the tap is indeed from a foe, which would warrant self-protection.

We have provided modelling-informed analysis of PTSD pathogenesis and related sequelae of traumatic stress and their relationship to diagnostic criteria in terms of vulnerability and resilience. Our key hypothesis is that the balance of plasticity-enhancement and plasticity-decay for aversive events is greater in the vulnerable. Below, we provide several novel auxiliary hypotheses that can be tested in controlled experiments.

Our analysis also provides a way to understand learning-based treatment mechanisms and their relation to (e.g.) monoaminergic interventions and substance abuse. Pharmacological interventions could enhance psychotherapies which support learning contextual disambiguation, through allocating attention to evidence for benign causes of current events, including aversive ones. This is a putative mechanism of action of entactogen-assisted psychotherapy. The combination of psychotherapy and psychopharmacology may be indispensable in some cases (e.g. due to TBI-related structural damage). At the same time, our account is consistent with psychotherapy providing sufficient treatment for trauma in many cases. ELEs fed back that understanding the interconnected aspects of our analysis can help mitigate self-stigma and shame (see Box 1, ‘ELE Feedback’), though establishing detailed mechanisms whereby social and therapy factors can help requires further work.

For onset prevention, the model suggests that arbitrary aspects of the environment during the post-traumatic period could hinder or induce impairment, depending on how strongly and/or frequently sensory cues mobilise a defensive response. For example, immediate debriefing following a traumatic experience could unduly remobilise a defensive response and strengthen memory of the harmful context. Social factors (e.g. support groups, transition homes) would have the potential to counterbalance adverse cued responses by strengthening the inference of a present harmless context and reducing trauma-adjacent emotional arousal, e.g. perception of powerlessness. Overall, our analysis lends support to biopsychosocial approaches to diagnosing and treating PTSD, or preventing its onset.

### Conclusions

Broadly speaking, investigating the indirect causes of PTSD in longitudinal studies remains important. At the same time, it is becoming increasingly possible to narrow investigations to more direct mechanisms. We take a step in this direction with our key hypothesis, that the balance of plasticity-enhancement and plasticity-decay for aversive events is greater in the vulnerable. Based on our findings, we expect that if genetic, developmental, and social factors (and their combination) are discovered that shift the neuroplasticity balance in the manner described, it will correspond to longitudinal studies of increased probability of PTSD.

More specifically, our analysis and modelling offer four hypotheses with testable predictions for immediate future work involving individuals with PTSD-like syndromes. First, our work predicts that in a psychophysical visual target/distractor paradigm that includes focal and peripheral sensing, cued traumatic recall in those with PTSD will attenuate sensory evidence accumulation at the periphery more than any attenuation occurring in focal regions; and with a greater differential than negatively valenced recall in unimpaired individuals. Relatedly, second, in a reality testing paradigm that measures stimulus congruency with mental imagery and confidence of reality judgments [91], traumatic recall in those with PTSD cued by a focal visual stimulus will show congruency and confident reality judgments even when incongruent peripheral cues are presented. Third, a psychophysical test of the exaggerated startle response (ESR) is predicted to reveal a lower differential between positively and negatively valenced contexts paired with the same probe stimulus in those with PTSD, as compared to unimpaired individuals [90]. Fourth, the active inference account presented here predicts that those with PTSD will show high precision in contextual modulation of trauma-associated cues; this contrasts with DRT, which predicts low top-down precision, allowing the theories to be experimentally distinguished [36].

Future work on modelling will explore the neuronal dynamics involved in *hierarchical prior dependent trauma re-experiencing*. According to our approach, this process can be linked not only to flashback and related phenomenology but also potentially to re-traumatisation. Formally, we understand re-traumatisation as the endogenous provision of pseudo-traumatic data that further amplifies the core mechanisms of the recurrent ABC model.

It remains an open question how maladaptive traumatic learning corresponds to ACh and NA. Yet the behaviour and precise neurobiology of these neuromodulators involved in sensory attenuation beyond initial learning consequently helped behavioural correspondence in a recent psychopharmacology study [97]. This is consistent with our simulations and supports the systematic mechanistic understanding presented here.

Overall, the present contribution comports with diverse empirical findings, suggesting a promising future route from basic to translational and clinical research on PTSD and related disorders. Further iterations of the modelling will aim to help identify more precise post-traumatic intervention targets. In turn, we hope this work will ultimately lead to increased prevention and treatment success for trauma-related mental health conditions.

#### Box 1

**Expert by Lived Experience (ELE) feedback**

*Seven ELEs individually reviewed this work and provided feedback, summarized here. Each point contains quotations from a single ELE (distinct from order listed in acknowledgments), reproduced verbatim apart from minor grammatical adjustments. Quotations have been re-ordered and grouped under thematic subheadings. Emphasis added in bold to highlight key takeaways*.

**Ability of the analysis to capture lived experience phenomena**

- Actually **made me feel understood** in a strange way. It’s like—my brain is doing what it thinks it should, assigning too much weight to the wrong signals. That finally explains why I feel like I’m always overreacting, even when I rationally know I’m safe.
- **The science gave me language for things I’ve sensed for a long time but didn’t know how to explain.** As someone with a background in military service and an interest in how the brain works, I really appreciated that this model goes beyond the old fear-conditioning stuff.
- This is **the first time I felt truly seen by a model**. When you described the brain clinging to old threat beliefs, **I felt like you were describing my daily life**. PTSD has always felt like being stuck in a story my body won’t stop telling—even when everything around me says I’m safe. This is the first time a model actually made sense of why I’m still reacting the way I do.
- **For me,** PTSD has never been about fear so much as **being on guard all the time**. It’s not just a reaction, it’s a posture. I’m always scanning, always planning escape routes, trying to control everything. That made total sense when I was in a dangerous environment—but now, it’s exhausting and doesn’t fit my life anymore. **The way you framed this** as ‘harm expectation’ and defensive mobilization **really captured that**.
- [The] focus on persistent hyper-alertness **struck a chord with me**, [it is] **on the mark based on my personal experience**. [The] concept of impaired belief-updating makes sense to me and could explain why all the reassurance I’ve received hasn’t helped. [How the analysis] describes flashbacks makes sense. [The] concept of self-reinforcing trauma cues makes a lot of sense, and **reflects my personal experience**.
- **This model really resonated** because it acknowledges that trauma doesn’t happen in a vacuum. My harm expectations are shaped by racism, cultural identity, and systemic violence. The concept of moral injury is huge for me. It’s not just about what happened, but how it conflicts with who I am, my values, and how I see the world. That’s often left out of clinical models.
- [The analysis is] **enlightening and re-assuring from a human perspective**. [It is] grounding and empowering, in that it helps develop a better sense of situational awareness and sense of self [which] improves confidence in self-efficacy. [It is] presented in a highly cogent and intuitive manner with clarity and nuance. [It] enhances understanding, meaning and clinical value in its broadest sense.

**Potential of the analysis to help transform the self-perception of affected individuals**

- The [analysis] **helps me trust that there’s a logic behind my symptoms**, even if it’s gone sideways. [The analysis] explains what my brain is doing, not just that it’s doing something weird.
- **I’ve always been ashamed** of the triggers I have. I feel like [the analysis] kind of

**sheds a new light on it**. Normally I just feel like the odd man out for how I feel.

- I appreciate that [the analysis] **doesn’t blame the individual**.
- [The analysis] can be of tremendous help in **destigmatising illness, including mitigating self-stigma and shame**.

**Suggestions to inform co-production of further research**

- [Include] more voices—especially from survivors, veterans, and marginalized communities. Emphasize how relational, cultural, and societal factors shape belief systems and recovery.
- [Include] more veteran voices.
- I’m really curious how this could translate into therapy—especially in ways that are relational and culturally aware.
- I’d be really interested in treatments that help recalibrate this [‘vicious circle’] process, rather than just manage the fallout.
- Keep making room for those of us whose trauma shows up more like hyper-preparation.
- Consider more directly how [institutions, cultures, and industries] can reinforce avoidance, over-medicalization, or mislabelling of trauma [which can] block or distort the healing process.

## Acknowledgements

The authors thank the Experts by Lived Experience (ELEs) who individually reviewed this work and provided feedback, including Rory Burke, Cody Cerne, and those who requested anonymity. AL received research funding support from the Open University School of Computing & Communications, and was hosted during manuscript editing by the Konrad Lorenz Institute for Evolution & Cognition Research. KF is supported by funding from the Wellcome Trust (Ref: 226793/Z/22/Z). The Functional Imaging Laboratory is funded by a Wellcome Discovery Grant. Beyond these disclosures, the authors received no specific funding for this work.

## Supplementary information

Simulations were performed in Matlab using a standard routine, spm_MDP_VB_X.m, included with the SPM software (http://www.fil.ion.ucl.ac.uk/spm/). Code available upon reasonable request (will be included in version of record). No Generative AI was used in the creation of this manuscript.

## References

1. Linson A, Parr T, Friston KJ. Active inference, stressors, and psychological trauma: A neuroethological model of (mal)adaptive explore-exploit dynamics in ecological context. Behav Brain Res. 2020 Feb 17;380:112421. doi:10.1016/j.bbr.2019.112421

2. Miles SR, Hale WJ, Mintz J, Wachen JS, Litz BT, Dondanville KA, et al. Hyperarousal symptoms linger after successful PTSD treatment in active duty military. Psychol Trauma Theory Res Pract Policy. 2023 Nov;15(8):1398–405. doi:10.1037/tra0001292

3. Cooke DF, Graziano MSA. Sensorimotor Integration in the Precentral Gyrus: Polysensory Neurons and Defensive Movements. J Neurophysiol. 2004 Apr 1;91(4):1648–60. doi:10.1152/jn.00955.2003

4. Cooke DF, Graziano MSA. Super-Flinchers and Nerves of Steel: Defensive Movements Altered by Chemical Manipulation of a Cortical Motor Area. Neuron. 2004 Aug 19;43(4):585–93. doi:10.1016/j.neuron.2004.07.029

5. Cisler JM, Dunsmoor JE, Fonzo GA, Nemeroff CB. Latent-state and model-based learning in PTSD. Trends Neurosci. 2024 Feb 1;47(2):150–62. doi:10.1016/j.tins.2023.12.002

6. Tsuda B, Pate SC, Tye KM, Siegelmann HT, Sejnowski TJ. Neuromodulators generate multiple context-relevant behaviors in a recurrent neural network by shifting activity flows in hyperchannels. bioRxiv. 2024 Jan 1;2021.05.31.446462. doi:10.1101/2021.05.31.446462

7. Mobbs D, Hagan CC, Dalgleish T, Silston B, Prévost C. The ecology of human fear: survival optimization and the nervous system. Front Neurosci. 2015;9. doi:10.3389/fnins.2015.00055

8. Maren S, Phan KL, Liberzon I. The contextual brain: implications for fear conditioning, extinction and psychopathology. Nat Rev Neurosci. 2013 Jun 1;14(6):417–28. doi:10.1038/nrn3492

9. VanElzakker MB, Kathryn Dahlgren M, Caroline Davis F, Dubois S, Shin LM. From Pavlov to PTSD: The extinction of conditioned fear in rodents, humans, and anxiety disorders. Extinction. 2014 Sep 1;113:3–18. doi:10.1016/j.nlm.2013.11.014

10. Bouyeure A, Pacheco D, Fellner MC, Jacob G, Kobelt M, Rose J, et al. Distinct representational properties of cues and contexts shape fear learning and extinction. 2025 Feb 27. doi:10.7554/elife.105126.1

11. Moutoussis M, Shahar N, Hauser TU, Dolan RJ. Computation in Psychotherapy, or How Computational Psychiatry Can Aid Learning-Based Psychological Therapies. Comput Psychiatry. 2018. doi:10.1162/CPSY_a_00014

12. Linson A, Friston K. Reframing PTSD for computational psychiatry with the active inference framework. Cognit Neuropsychiatry. 2019;24(5):347–68. doi:10.1080/13546805.2019.1665994

13. Seriès P. Post-traumatic stress disorder as a disorder of prediction. Nat Neurosci. 2019 Mar 1;22(3):334–6. doi:10.1038/s41593-019-0345-z

14. Seriès P, Veerapa E, Jardri R. Can computational models help elucidate the link between complex trauma and hallucinations? Hallucinations Neurobiol Patient Exp. 2024 Mar 1;265:66–73. doi:10.1016/j.schres.2023.05.003

15. Van der Kolk B, Spinazzola J, Blaustein M, Hopper J, Hopper E, Korn D, et al. A randomized clinical trial of EMDR, fluoxetine and pill placebo in the treatment of PTSD: Treatment effects and long-term maintenance. J Clin Psychiatry. 2007;68(1):37–46. doi:10.4088/jcp.v68n0105

16. Kaye AP, Rao MG, Kwan AC, Ressler KJ, Krystal JH. A computational model for learning from repeated traumatic experiences under uncertainty. Cogn Affect Behav Neurosci. 2023 Jun 1;23(3):894–904. doi:10.3758/s13415-023-01085-5

17. Harnett NG, Fleming LL, Clancy KJ, Ressler KJ, Rosso IM. Affective Visual Circuit Dysfunction in Trauma and Stress-Related Disorders. Biol Psychiatry. 2025 Feb 15;97(4):405–16. doi:10.1016/j.biopsych.2024.07.003

18. Lange RD, Chattoraj A, Beck JM, Yates JL, Haefner RM. A confirmation bias in perceptual decision-making due to hierarchical approximate inference. PLOS Comput Biol. 2021 Nov 29;17(11):e1009517. doi:10.1371/journal.pcbi.1009517

19. Shohamy D, Adcock RA. Dopamine and adaptive memory. Trends Cogn Sci. 2010 Oct 1;14(10):464–72. doi:10.1016/j.tics.2010.08.002

20. Tomić I, Bays PM. A dynamic neural resource model bridges sensory and working memory. Salinas E, Gold JI, editors. eLife. 2024 May 3;12:RP91034. doi:10.7554/eLife.91034

21. Cotton K, Ricker TJ. Examining the relationship between working memory consolidation and long-term consolidation. Psychon Bull Rev. 2022 Oct 1;29(5):1625–48. doi:10.3758/s13423-022-02084-2

22. Friston K, FitzGerald T, Rigoli F, Schwartenbeck P, Pezzulo G. Active inference: a process theory. Neural Comput. 2017;29(1):1–49. doi:10.1162/NECO_a_00912

23. Smith R, Friston KJ, Whyte CJ. A step-by-step tutorial on active inference and its application to empirical data. J Math Psychol. 2022 Apr 1;107:102632. doi:10.1016/j.jmp.2021.102632

24. Horner AJ, Burgess N. The associative structure of memory for multi-element events. J Exp Psychol Gen. 2013;142(4):1370–83. doi:10.1037/a0033626

25. Horner AJ, Bisby JA, Bush D, Lin WJ, Burgess N. Evidence for holistic episodic recollection via hippocampal pattern completion. Nat Commun. 2015 Jul 2;6(1):7462. doi:10.1038/ncomms8462

26. Koolschijn RS, Shpektor A, Clarke WT, Ip IB, Dupret D, Emir UE, et al. Memory recall involves a transient break in excitatory-inhibitory balance. Irish M, Baker CI, Mullins P, Hutchinson B, editors. eLife. 2021 Oct 8;10:e70071. doi:10.7554/eLife.70071

27. Holton E, Braun L, Thompson JAF, Grohn J, Summerfield C. Humans and neural networks show similar patterns of transfer and interference during continual learning [Internet]. OSF; 2025 [cited 2025 Feb 28]. Available from: https://osf.io/98ksw_v1 doi:10.31234/osf.io/98ksw_v1

28. Park J, Holmes CD, Snyder LH. Compositional architecture: Orthogonal neural codes for task context and spatial memory in prefrontal cortex. bioRxiv. 2025 Jan 1;2025.02.25.640211. doi:10.1101/2025.02.25.640211

29. Wills TJ, Lever C, Cacucci F, Burgess N, O’Keefe J. Attractor Dynamics in the Hippocampal Representation of the Local Environment. Science. 2005 May 6;308(5723):873–6. doi:10.1126/science.1108905

30. McKenzie S, Frank AJ, Kinsky NR, Porter B, Rivière PD, Eichenbaum H. Hippocampal Representation of Related and Opposing Memories Develop within Distinct, Hierarchically Organized Neural Schemas. Neuron. 2014 Jul 2;83(1):202–15. doi:10.1016/j.neuron.2014.05.019

31. Ólafsdóttir HF, Carpenter F, Barry C. Task Demands Predict a Dynamic Switch in the Content of Awake Hippocampal Replay. Neuron. 2017 Nov 15;96(4):925–935.e6. doi:10.1016/j.neuron.2017.09.035

32. Favila SE, Chanales AJH, Kuhl BA. Experience-dependent hippocampal pattern differentiation prevents interference during subsequent learning. Nat Commun. 2016 Apr 6;7(1):11066. doi:10.1038/ncomms11066

33. Zotow E, Bisby JA, Burgess N. Behavioral evidence for pattern separation in human episodic memory. Learn Mem. 2020 Aug 1;27(8):301–9. doi:10.1101/lm.051821.120

34. Friston K, FitzGerald T, Rigoli F, Schwartenbeck P, O’Doherty J, Pezzulo G. Active inference and learning. Neurosci Biobehav Rev. 2016 Sep 1;68:862–79. doi:10.1016/j.neubiorev.2016.06.022

35. Friston K, Buzsáki G. The Functional Anatomy of Time: What and When in the Brain. Trends Cogn Sci. 2016 Jul 1;20(7):500–11. doi:10.1016/j.tics.2016.05.001

36. Brewin CR, Burgess N. Contextualisation in the revised dual representation theory of PTSD: A response to Pearson and colleagues. J Behav Ther Exp Psychiatry. 2014 Mar 1;45(1):217–9. doi:10.1016/j.jbtep.2013.07.011

37. Sarto-Jackson I. The Making and Breaking of Minds: How Social Interactions Shape the Human Mind. Vernon Press; 2022.

38. Heinbockel H, Wagner AD, Schwabe L. Post-retrieval stress impairs subsequent memory depending on hippocampal memory trace reinstatement during reactivation. Sci Adv. 10(18):eadm7504. doi:10.1126/sciadv.adm7504

39. Perl O, Duek O, Kulkarni KR, Gordon C, Krystal JH, Levy I, et al. Neural patterns differentiate traumatic from sad autobiographical memories in PTSD. Nat Neurosci. 2023 Dec 1;26(12):2226–36. doi:10.1038/s41593-023-01483-5

40. Mellman TA, Bustamante V, Fins AI, Pigeon WR, Nolan B. REM Sleep and the Early Development of Posttraumatic Stress Disorder. Am J Psychiatry. 2002 Oct 1;159(10):1696–701. doi:10.1176/appi.ajp.159.10.1696

41. Mellman TA, Kulick-Bell R, Ashlock LE, Nolan B. Sleep events among veterans with combat-related posttraumatic stress disorder. Am J Psychiatry. 1995;152(1):110–5. doi:10.1176/ajp.152.1.110

42. Chang H, Tang W, Wulf AM, Nyasulu T, Wolf ME, Fernandez-Ruiz A, et al. Sleep microstructure organizes memory replay. Nature. 2025 Jan 1;637(8048):1161–9. doi:10.1038/s41586-024-08340-w

43. Bollmann L, Baracskay P, Stella F, Csicsvari J. Sleep stages antagonistically modulate reactivation drift. Neuron. 2025. doi:10.1016/j.neuron.2025.02.025

44. Balouek JA, Mclain CA, Minerva AR, Rashford RL, Bennett SN, Rogers FD, et al. Reactivation of Early-Life Stress-Sensitive Neuronal Ensembles Contributes to Lifelong Stress Hypersensitivity. J Neurosci. 2023 Aug 23;43(34):5996–6009. doi:10.1523/JNEUROSCI.0016-23.2023

45. Hobson JA, Friston KJ. Waking and dreaming consciousness: Neurobiological and functional considerations. Prog Neurobiol. 2012 Jul 1;98(1):82–98. doi:10.1016/j.pneurobio.2012.05.003

46. Geva N, Deitch D, Rubin A, Ziv Y. Time and experience differentially affect distinct aspects of hippocampal representational drift. Neuron. 2023 Aug 2;111(15):2357–2366.e5. doi:10.1016/j.neuron.2023.05.005

47. McNamara P, Wildman WJ, Hodulik G, Rohr D. A neurocomputational theory of nightmares: the role of formal properties of nightmare images. SLEEP Adv. 2021 Mar 1;2(1):zpab009. doi:10.1093/sleepadvances/zpab009

48. Garcia-Rill E. Disorders of the reticular activating system. Med Hypotheses. 1997 Nov 1;49(5):379–87. doi:10.1016/S0306-9877(97)90083-9

49. Hobson JA, Gott JA, Friston KJ. Minds and Brains, Sleep and Psychiatry. Psychiatr Res Clin Pract. 2021 Mar 1;3(1):12–28. doi:10.1176/appi.prcp.20200023

50. Mendoza Alvarez M, Balthasar Y, Verbraecken J, Claes L, van Someren E, van Marle HJF, et al. Systematic review: REM sleep, dysphoric dreams and nightmares as transdiagnostic features of psychiatric disorders with emotion dysregulation - Clinical implications. Sleep Med. 2025 Mar 1;127:1–15. doi:10.1016/j.sleep.2024.12.037

51. Krueger J, Disney AA. Structure and function of dual-source cholinergic modulation in early vision. J Comp Neurol. 2019 Feb 15;527(3):738–50. doi:10.1002/cne.24590

52. Garcia-Rill E. Chapter 5 - Posttraumatic stress and anxiety, the role of arousal. In: Garcia-Rill E, editor. Arousal in Neurological and Psychiatric Diseases. Academic Press; 2019. p. 67–81. doi:10.1016/B978-0-12-817992-5.00005-2

53. Gott JA, Stücker S, Kanske P, Haaker J, Dresler M. Acetylcholine and metacognition during sleep. Conscious Cogn. 2024 Jan 1;117:103608. doi:10.1016/j.concog.2023.103608

54. Ben-Zion Z, Simon AJ, Rosenblatt M, Korem N, Duek O, Liberzon I, et al. Connectome-Based Predictive Modeling of PTSD Development Among Recent Trauma Survivors. JAMA Netw Open. 2025 Mar 10;8(3):e250331. doi:10.1001/jamanetworkopen.2025.0331

55. Zhou L, Liu MZ, Li Q, Deng J, Mu D, Sun YG. Organization of Functional Long-Range Circuits Controlling the Activity of Serotonergic Neurons in the Dorsal Raphe Nucleus. Cell Rep. 2017 Mar 21;18(12):3018–32. doi:10.1016/j.celrep.2017.02.077

56. Michels L, Schulte-Vels T, Schick M, O’Gorman RL, Zeffiro T, Hasler G, et al. Prefrontal GABA and glutathione imbalance in posttraumatic stress disorder: Preliminary findings. Psychiatry Res Neuroimaging. 2014 Dec 30;224(3):288–95. doi:10.1016/j.pscychresns.2014.09.007

57. Nyberg L, Karalija N, Salami A, Andersson M, Wåhlin A, Kaboovand N, et al. Dopamine D2 receptor availability is linked to hippocampal–caudate functional connectivity and episodic memory. Proc Natl Acad Sci. 2016 Jul 12;113(28):7918–23. doi:10.1073/pnas.1606309113

58. Schott BH, Seidenbecher CI, Fenker DB, Lauer CJ, Bunzeck N, Bernstein HG, et al. The Dopaminergic Midbrain Participates in Human Episodic Memory Formation: Evidence from Genetic Imaging. J Neurosci. 2006 Feb 1;26(5):1407. doi:10.1523/JNEUROSCI.3463-05.2006

59. Bentley P, Vuilleumier P, Thiel CM, Driver J, Dolan RJ. Cholinergic enhancement modulates neural correlates of selective attention and emotional processing. NeuroImage. 2003 Sep 1;20(1):58–70. doi:10.1016/S1053-8119(03)00302-1

60. Picciotto MR, Higley MJ, Mineur YS. Acetylcholine as a Neuromodulator: Cholinergic Signaling Shapes Nervous System Function and Behavior. Neuron. 2012 Oct 4;76(1):116–29. doi:10.1016/j.neuron.2012.08.036

61. Naegeli C, Zeffiro T, Piccirelli M, Jaillard A, Weilenmann A, Hassanpour K, et al. Locus Coeruleus Activity Mediates Hyperresponsiveness in Posttraumatic Stress Disorder. Biol Psychiatry. 2018 Feb 1;83(3):254–62. doi:10.1016/j.biopsych.2017.08.021

62. Beauchamp MS, Yasar NE, Frye RE, Ro T. Touch, sound and vision in human superior temporal sulcus. NeuroImage. 2008 Jul 1;41(3):1011–20. doi:10.1016/j.neuroimage.2008.03.015

63. Butter CM, Buchtel HA, Santucci R. Spatial attentional shifts: Further evidence for the role of polysensory mechanisms using visual and tactile stimuli. Neuropsychologia. 1989 Jan 1;27(10):1231–40. doi:10.1016/0028-3932(89)90035-3

64. Wald A. An Essentially Complete Class of Admissible Decision Functions. Ann Math Stat. 1947 Dec;18(4):549–55. doi:10.1214/aoms/1177730345

65. Brown LD. A Complete Class Theorem for Statistical Problems with Finite Sample Spaces. Ann Stat. 1981 Nov;9(6):1289–300. doi:10.1214/aos/1176345645

66. Reddy L, Kanwisher N. Category Selectivity in the Ventral Visual Pathway Confers Robustness to Clutter and Diverted Attention. Curr Biol. 2007 Dec 4;17(23):2067–72. doi:10.1016/j.cub.2007.10.043

67. Mueller-Pfeiffer C, Schick M, Schulte-Vels T, O’Gorman R, Michels L, Martin-Soelch C, et al. Atypical visual processing in posttraumatic stress disorder. NeuroImage Clin. 2013 Jan 1;3:531–8. doi:10.1016/j.nicl.2013.08.009

68. Da Costa L, Parr T, Sajid N, Veselic S, Neacsu V, Friston K. Active inference on discrete state-spaces: A synthesis. J Math Psychol. 2020 Dec 1;99:102447. doi:10.1016/j.jmp.2020.102447

69. Parr T, Da Costa L, Friston K. Markov blankets, information geometry and stochastic thermodynamics. Philos Trans R Soc Math Phys Eng Sci. 2019 Dec 23;378(2164):20190159. doi:10.1098/rsta.2019.0159

70. Da Costa L, Parr T, Sengupta B, Friston K. Neural Dynamics under Active Inference: Plausibility and Efficiency of Information Processing. Entropy. 2021;23(4). doi:10.3390/e23040454

71. Wurtz RH, McAlonan K, Cavanaugh J, Berman RA. Thalamic pathways for active vision. Trends Cogn Sci. 2011 Apr 1;15(4):177–84. doi:10.1016/j.tics.2011.02.004

72. Parr T, Friston KJ. The active construction of the visual world. Neuropsychologia. 2017 Sep 1;104:92–101. doi:10.1016/j.neuropsychologia.2017.08.003

73. Mirza MB, Adams RA, Mathys C, Friston KJ. Human visual exploration reduces uncertainty about the sensed world. PLOS ONE. 2018 Jan 5;13(1):e0190429. doi:10.1371/journal.pone.0190429

74. Rao RPN, Gklezakos DC, Sathish V. Active Predictive Coding: A Unifying Neural Model for Active Perception, Compositional Learning, and Hierarchical Planning. Neural Comput. 2023 Dec 12;36(1):1–32. doi:10.1162/neco_a_01627

75. Dijkstra N, Zeidman P, Ondobaka S, van Gerven MAJ, Friston K. Distinct Top-down and Bottom-up Brain Connectivity During Visual Perception and Imagery. Sci Rep. 2017 Jul 18;7(1):5677. doi:10.1038/s41598-017-05888-8

76. Albers AM, Kok P, Toni I, Dijkerman HC, de Lange FP. Shared Representations for Working Memory and Mental Imagery in Early Visual Cortex. Curr Biol. 2013 Aug 5;23(15):1427–31. doi:10.1016/j.cub.2013.05.065

77. Schwartz S, Vuilleumier P, Hutton C, Maravita A, Dolan RJ, Driver J. Attentional Load and Sensory Competition in Human Vision: Modulation of fMRI Responses by Load at Fixation during Task-irrelevant Stimulation in the Peripheral Visual Field. Cereb Cortex. 2005 Jun 1;15(6):770–86. doi:10.1093/cercor/bhh178

78. Stolte M, Bahrami B, Lavie N. High perceptual load leads to both reduced gain and broader orientation tuning. J Vis. 2014 Mar 7;14(3):9–9. doi:10.1167/14.3.9

79. Torralbo A, Kelley TA, Rees G, Lavie N. Attention induced neural response trade-off in retinotopic cortex under load. Sci Rep. 2016 Sep 14;6(1):33041. doi:10.1038/srep33041

80. Clancy K, Ding M, Bernat E, Schmidt NB, Li W. Restless ‘rest’: intrinsic sensory hyperactivity and disinhibition in post-traumatic stress disorder. Brain. 2017 Jul 1;140(7):2041–50. doi:10.1093/brain/awx116

81. Harrison SA, Tong F. Decoding reveals the contents of visual working memory in early visual areas. Nature. 2009 Apr 1;458(7238):632–5. doi:10.1038/nature07832

82. Teng C, Kravitz DJ. Visual working memory directly alters perception. Nat Hum Behav. 2019 Aug 1;3(8):827–36. doi:10.1038/s41562-019-0640-4

83. S. A. Brandt, L. W. Stark. Spontaneous Eye Movements During Visual Imagery Reflect the Content of the Visual Scene. J Cogn Neurosci. 1997 Jan;9(1):27–38. doi:10.1162/jocn.1997.9.1.27

84. Spivey MJ, Geng JJ. Oculomotor mechanisms activated by imagery and memory: eye movements to absent objects. Psychol Res. 2001 Nov 1;65(4):235–41. doi:10.1007/s004260100059

85. Andrade J, Kavanagh D, Baddeley A. Eye-movements and visual imagery: A working memory approach to the treatment of post-traumatic stress disorder. Br J Clin Psychol. 1997 May 1;36(2):209–23. doi:10.1111/j.2044-8260.1997.tb01408.x

86. Itti L, Baldi P. Bayesian surprise attracts human attention. Vis Atten Psychophys Electrophysiol Neuroimaging. 2009 Jun 2;49(10):1295–306. doi:10.1016/j.visres.2008.09.007

87. Sun Y, Gomez F, Schmidhuber J. Planning to Be Surprised: Optimal Bayesian Exploration in Dynamic Environments. In: Schmidhuber J, Thórisson KR, Looks M, editors. Artificial General Intelligence. Berlin, Heidelberg: Springer Berlin Heidelberg; 2011. p. 41–51. doi:10.1007/978-3-642-22887-2_5

88. Parr T, Friston KJ. Working memory, attention, and salience in active inference. Sci Rep. 2017 Nov 7;7(1):14678. doi:10.1038/s41598-017-15249-0

89. Parr T, Friston KJ. Attention or salience? Atten Percept. 2019 Oct 1;29:1–5. doi:10.1016/j.copsyc.2018.10.006

90. Bradley MM, Cuthbert BN, Lang PJ. Pictures as prepulse: Attention and emotion in startle modification. Psychophysiology. 1993 Sep 1;30(5):541–5. doi:10.1111/j.1469-8986.1993.tb02079.x

91. Dijkstra N, von Rein T, Kok P, Fleming SM. A neural basis for distinguishing imagination from reality. Neuron. 2025. doi:10.1016/j.neuron.2025.05.015

92. Monsalve-Mercado MM, Stine GM, Shadlen MN, Miller KD. The geometry of the neural state space of decisions. bioRxiv. 2025 Jan 1;2025.01.24.634806. doi:10.1101/2025.01.24.634806

93. Mirza MB, Adams RA, Friston K, Parr T. Introducing a Bayesian model of selective attention based on active inference. Sci Rep. 2019 Sep 26;9(1):13915. doi:10.1038/s41598-019-50138-8

94. Khona M, Fiete IR. Attractor and integrator networks in the brain. Nat Rev Neurosci. 2022 Dec 1;23(12):744–66. doi:10.1038/s41583-022-00642-0

95. Chiarotti F, Viglione A, Giuliani A, Branchi I. Citalopram amplifies the influence of living conditions on mood in depressed patients enrolled in the STAR*D study. Transl Psychiatry. 2017 Mar;7(3):e1066–e1066. doi:10.1038/tp.2017.35

96. Carhart-Harris RL, Chandaria S, Erritzoe DE, Gazzaley A, Girn M, Kettner H, et al. Canalization and plasticity in psychopathology. Neuropharmacology. 2023 Mar 15;226:109398. doi:10.1016/j.neuropharm.2022.109398

97. Sogo K, Sogo M, Okawa Y. Centrally acting anticholinergic drug trihexyphenidyl is highly effective in reducing nightmares associated with post-traumatic stress disorder. Brain Behav. 2021 Jun 1;11(6):e02147. doi:10.1002/brb3.2147

